# The genomic landscape of intrinsic and acquired resistance to cyclin-dependent kinase 4/6 inhibitors in patients with hormone receptor positive metastatic breast cancer

**DOI:** 10.1101/857839

**Authors:** Seth A. Wander, Ofir Cohen, Xueqian Gong, Gabriela N. Johnson, Jorge Buendia-Buendia, Maxwell R. Lloyd, Dewey Kim, Flora Luo, Pingping Mao, Karla Helvie, Kailey J. Kowalski, Utthara Nayar, Adrienne G. Waks, Stephen Parsons, Ricardo Martinez, Lacey M. Litchfield, Xiang S. Ye, Chun Ping Yu, Valerie M. Jansen, John R. Stille, Patricia S. Smith, Gerard J. Oakley, Quincy Chu, Gerald Batist, Melissa Hughes, Jill D. Kremer, Levi A. Garraway, Eric P. Winer, Sara M. Tolaney, Nancy U. Lin, Sean Buchanan, Nikhil Wagle

## Abstract

Clinical resistance mechanisms to CDK4/6 inhibitors in HR+ breast cancer have not been clearly defined. Whole exome sequencing of 59 tumors with CDK4/6i exposure revealed multiple candidate resistance mechanisms including *RB1* loss, activating alterations in *AKT1*, *RAS*, *AURKA*, *CCNE2, ERBB2,* and *FGFR2,* and loss of ER expression. *In vitro* experiments confirmed that these alterations conferred CDK4/6i resistance. Cancer cells cultured to resistance with CDK4/6i also acquired *RB1*, *KRAS*, *AURKA*, or *CCNE2* alterations, which conferred sensitivity to AURKA, ERK, or CHEK1 inhibition. Besides inactivation of RB1, which accounts for ∼5% of resistance, seven of these mechanisms have not been previously identified as clinical mediators of resistance to CDK4/6 inhibitors in patients. Three of these—RAS activation, AKT activation, and AURKA activation—have not to our knowledge been previously demonstrated preclinically. Together, these eight mechanisms were present in 80% of resistant tumors profiled and may define therapeutic opportunities in patients.

**Significance:** We identified eight distinct mechanisms of resistance to CDK4/6 inhibitors present in 80% of resistant tumors profiled. Most of these have a therapeutic strategy to overcome or prevent resistance in these tumors. Taken together, these findings have critical implications related to the potential utility of precision-based approaches to overcome resistance in many patients with HR+ MBC.

## Introduction

The cyclin-dependent kinase 4/6 inhibitors (CDK4/6i) have entered widespread use in both the first- and subsequent-line setting for patients with hormone-receptor positive (HR+), human epidermal growth factor receptor 2 negative (HER2-) metastatic breast cancer (MBC).^1, 2^ Their application has resulted in significant improvements in progression-free survival (PFS) and overall survival (OS) for treatment-naïve and previously treated patients in combination with anti-estrogens.^3–9^ Abemaciclib has shown efficacy as a single agent in endocrine-refractory disease, and has been approved for use as monotherapy in pre-treated patients with HR+/HER2-MBC.^10^ Despite these advances, HR+/HER2-MBC remains a significant cause of morbidity and mortality. Many patients demonstrate *de novo*, or intrinsic, resistance to these agents and, in those who respond, acquired resistance and disease progression is unfortunately inevitable.

We have limited insight into the molecular pathways governing resistance to CDK4/6i. Early development of these compounds indicated preferential efficacy in luminal/Rb-positive cell lines.^11^ Loss of Rb expression has been identified in cellular models cultured to resistance in CDK4/6i.^12^ Acquired *RB1* loss-of-function mutations were identified in circulating tumor DNA (ctDNA) from three patients following progression on CDK4/6i-based therapy.^13^ Analysis of ctDNA from patients treated on the PALOMA-3 trial, which explored palbociclib with fulvestrant versus fulvestrant alone in the second-line metastatic setting, demonstrated rare *RB1* mutations that were uniquely present in the group receiving palbociclib.^14^ *PIK3CA* and *ESR1* mutations were identified frequently on both arms of the study, and neither has been well established as a predictive biomarker.^14, 15^ Recent analysis of ctDNA and tumors from the MONARCH-2 study, exploring abemaciclib and fulvestrant in patients with prior progression on anti-estrogen therapy, suggested benefit from abemaciclib use regardless of *PIK3CA* or *ESR1* status, though the magnitude of benefit was larger in mutant patients.^16^ Despite lack of robust data supporting a role for PI3K, loss of the *PTEN* tumor suppressor was recently noted in tumor samples with progression on ribociclib, and was sufficient to promote resistance *in vitro*.^17^ Preclinically, PDK1, another PI3K pathway effector, emerged from a kinome-wide screen in HR+ cells as a potential mediator of resistance to CDK4/6i; targeting PDK1 or PI3K prompted resensitization to CDK4/6i.^18^

Preclinical studies have also implicated overexpression of CDK6 and cyclin E2 (CCNE2) in mediating resistance.^19, 20^ Increased expression of cyclin E1 (CCNE1) was associated with inferior response to palbociclib on PALOMA-3, while the expression of cyclin D1, RB1, and CDK4 failed to demonstrate any association.^21^ Targeted sequencing of tumor specimens from patients with HR+ MBC and CDK4/6i exposure suggested that regulation of CDK6 expression via the FAT1 tumor suppressor could provoke resistance^22^ and CDK6 expression may also be regulated via micro-RNA-dependent modulation of the TGF-B pathway, altering sensitivity to CDK4/6i *in vitro* and in patients.^23^

Prior work from our laboratory has implicated alterations in *ERBB2* and *FGFR2* in mediating resistance to CDK4/6i *in vitro* and in patients.^24, 25^ In addition, amplification of *FGFR1*, identified via sequencing of ctDNA from MONALEESA-2 (ribociclib and anti-estrogen in the first-line metastatic setting), correlated with reduced PFS and activation of FGFR1 provoked resistance *in vitro.*^26^

Here we explore the genomic landscape of resistance to CDK4/6i via whole exome sequencing of tumor biopsies. The landscape of resistance to CDK4/6i is heterogeneous, with multiple potential mediators including biallelic *RB1* disruption and activation of *AKT1*, *RAS*, *ERBB2*, *FGFR2*, Aurora Kinase A (*AURKA*), and *CCNE2*. Modification of HR+ breast cancer cells, via CRISPR-mediated knockout or lentiviral overexpression, corroborates the candidate mechanisms of resistance identified by tumor sequencing. Cells cultured to resistance in the presence of CDK4/6i demonstrate concordant alterations in RB1, AURKA, and CCNE2 expression along with RAS/ERK activation and demonstrate enhanced sensitivity to novel targeted therapies. In one patient with HR+/HER2-MBC that progressed on first-line CDK4/6i, AURKA inhibition provoked prolonged disease control in a phase I clinical trial. These results shed new light on the diverse landscape of genomic alterations that drive resistance to CDK4/6i in HR+/HER2-MBC and provide preclinical and translational rationale for novel strategies to circumvent and overcome resistance.

## Results

### The genomic landscape of intrinsic and acquired CDK4/6i resistance

We identified patients with HR+/HER2-MBC who were treated with CDK4/6i with or without an anti-estrogen and provided metastatic tumor biopsies as part of an IRB-approved tissue collection protocol.^27^ We classified samples as reflecting sensitivity, intrinsic resistance, or acquired resistance (Figure 1A). Sensitive biopsies were defined as baseline samples obtained within 120 days prior to, or up to a maximum of 31 days after, CDK4/6i initiation in a patient with subsequent clinical benefit (defined as radiographic response or stable disease >6 months). Biopsies reflecting intrinsic resistance were obtained within 120 days prior to or anytime after CDK4/6i initiation in patients without evidence of clinical benefit (defined as progression on the first interval restaging study or stable disease <6 months). Biopsies reflecting acquired resistance were obtained from patients who had experienced clinical benefit with CDK4/6i and had an available biopsy specimen within 31 days prior to progression or at any time thereafter.

**Figure 1.**
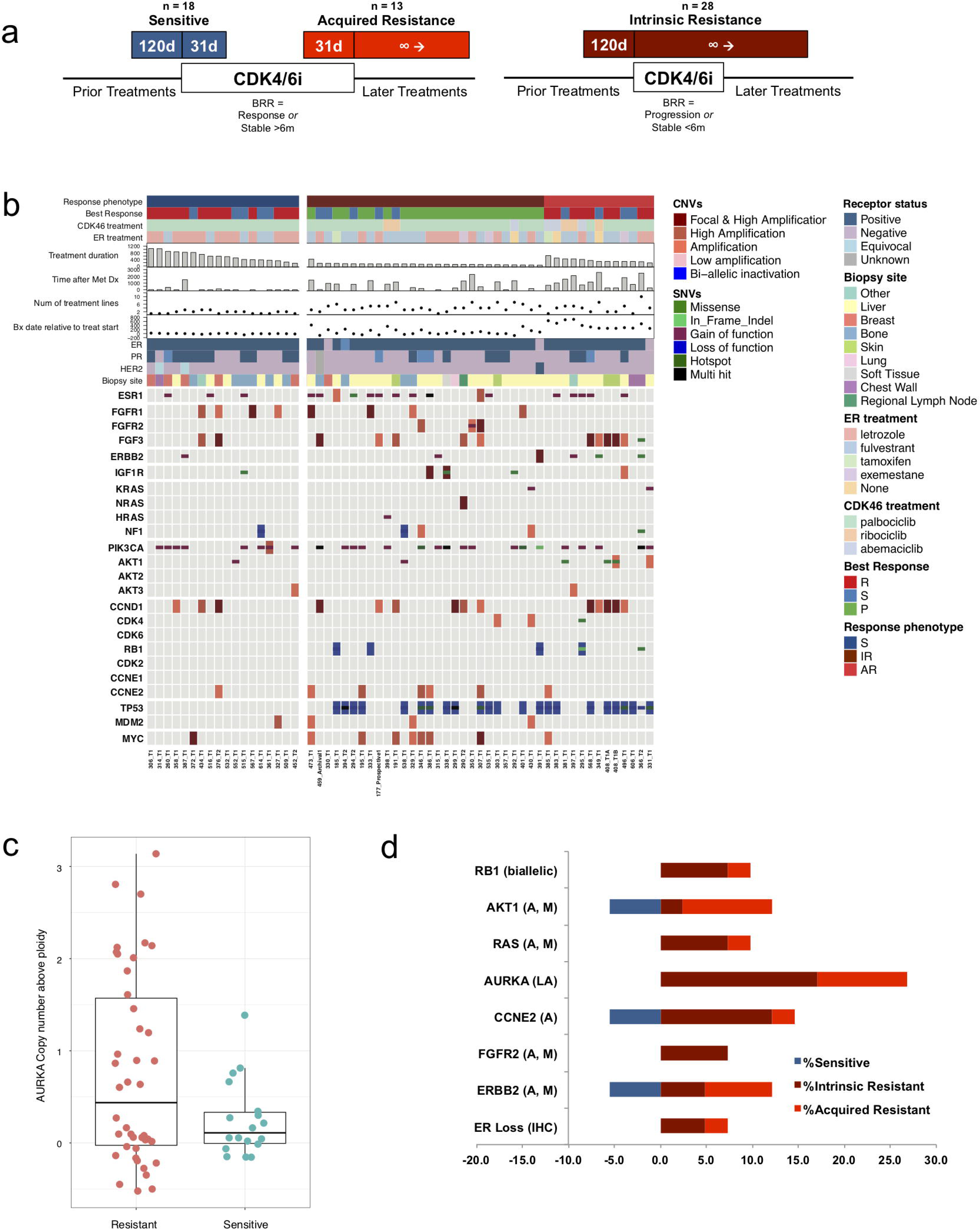
The genomic landscape of CDK4/6i resistance is heterogeneous, with multiple potential driver events. (a) Biopsy phenotypes were assigned as *sensitive, acquired resistance,* or *intrinsic resistance* based upon timing of the biopsy relative to CDK4/6i exposure (d - days), best radiographic response (BRR), and duration of treatment. Patients were categorized as experiencing clinical benefit on CDK4/6i if interval restaging demonstrated a response or disease stability for at least six months. (b) Mutational matrix (CoMut) depicting the genomic landscape of the CDK4/6i cohort (n = 59 biopsies, 58 patients). Copy number alterations and mutational events in select genes of interest are shown. Clinical parameters (shown at the top) include receptor status, anti-estrogen agent, CDK4/6 inhibitor, best radiographic response (P – progression, R – response, S – stable), biopsy phenotype (S – sensitive, IR – intrinsic resistance, AR – acquired resistance), treatment duration (days), biopsy timing relative to treatment initiation (days), time since metastatic diagnosis (days), and number of lines of prior treatment. (c) Phenotype distribution plot demonstrating a higher frequency of copy number amplifications in Aurora Kinase A (AURKA) among resistant biopsies (AR + IR, left) compared to sensitive biopsies (right, 0.0081, Welch test). (d) Bar plot visualization of mutational (M) and/or copy number alterations (A – amplification, LA – low amplification) in select genes. The proportional enrichment (fraction of samples demonstrating alteration) in sensitive biopsies (left, blue) and resistant biopsies (AR + IR, right, red) is included.

WES was successfully performed on 59 biopsies from 58 patients within the appropriate exposure window to be assigned a phenotype and with sufficient clinical data to define response (w). This included 18 sensitive biopsies, 28 intrinsic resistance biopsies, and 13 acquired resistance biopsies. The majority of patients (55, 94.8%) received standard combinations of an aromatase inhibitor or fulvestrant and a CDK4/6 inhibitor. 49 patients (84.5%) received a palbociclib-based regimen, including 28 patients (48.3%) with an aromatase inhibitor and 20 patients (34.5%) with fulvestrant. The mean duration of therapy was 316 days (range 43-1052). Patients received an average of 1.5 lines of therapy in the metastatic setting (range 0-7) and 30 patients (51.7%) had prior anti-estrogen exposure in the metastatic setting. Additional clinical parameters are described in Supplemental Table 2.

Whole exome sequencing of all 59 samples demonstrated a number of genomic alterations in genes implicated in HR+ breast cancer (*ESR1, PIK3CA, CCND1, FGFR1, TP53*) as well as additional cancer genes and putative resistance mediators (*RB1, ERBB2, FGFR2, AKT1, KRAS, HRAS, NRAS*, among others) (Figure 1B, Supplemental Table 3). Many of these alterations were enriched in resistant samples and not present or relatively infrequent in sensitive samples, suggesting they might be contributing to resistance (Figure 1B; Supplemental Figure 1; Supplemental Table 4). In addition to these genomic differences, three patients with resistant tumor biopsies demonstrated loss of ER expression in the metastatic drug-resistant tumor (measured by immunohistochemistry); all patients were known to be ER+ at the time of metastatic diagnosis.

While isolated amplification events were identified in a variety of cancer genes (Supplemental Table 4), amplification events in aurora kinase A were observed as occurring more frequently in resistant samples as compared with sensitive (0 in sensitive, 26.8% in resistant; 0.0081, Welch test) (Figure 1C). While only moderate magnitude AURKA amplifications were seen among the resistant tumors, in The Cancer Genome Atlas (TCGA) study, even low AURKA amplification in primary HR+ breast cancer samples resulted in a statistically significant increase in gene expression (Supplementary Figure 2), suggesting that the degree of AURKA amplification observed in the CDK4/6i-resistant cohort is likely to have a meaningful effect on gene expression and protein level.

Based on prior preclinical studies and known biology, we hypothesized that the following eight specific categories of alterations that were enriched in the resistant tumors were contributing to CDK4/6i resistance: biallelic disruption of *RB1*, activating mutation and/or amplification of *AKT1*, activating mutations in *KRAS/HRAS/NRAS*, activating mutations and/or amplification of *FGFR2*, activating mutations in *ERBB2*, amplification of CCNE2, amplification of AURKA, and loss of ER.

In total, 33 out of the 41 resistant biopsies (80.5%) had genomic alterations in at least one of these 8 potential resistance mechanisms, as compared to 3 of the 18 sensitive biopsies (Figure 1D, Supplemental Table 5). Consistent with prior reports, biallelic disruption in *RB1* was exclusively present in resistant samples and occurred in a minority of resistant biopsies (n=4/41, 9.8%). We identified diverse mechanisms of biallelic *RB1* disruption across the affected patients. In all examples, a single copy loss was noted in the presence of a point mutation, splice site alteration, or frameshift event in the second allele.

AKT1 alterations were identified in five resistant biopsies (n=5/41, 12.2%), including both mutational events and amplifications. A single sensitive biopsy also demonstrated an activating AKT1 alteration (n=1/18, 5.6%).

Diverse RAS-pathway activating events were observed in four CDK4/6i-resistant cases (n=4/41, 9.8%) including canonical activating mutations in *KRAS* G12D, a pathogenic mutation in KRAS Q61L,^28^ a mutation in *HRAS* K117N,^29^ and high focal amplification in *NRAS* (Figure 1B). There were no instances of RAS-altered tumors with a sensitive phenotype.

Amplification events in AURKA were identified in eleven resistant biopsies (n=11/41, 26.8%), including examples of both intrinsic and acquired resistance (n=7 and n=4, respectively). There were no sensitive biopsies with AURKA amplification.

There were six instances (n=6/41, 14.6%) of CCNE2 amplification identified across the resistant cohort (Figure 1B). A single sensitive biopsy with a CCNE2 alteration was identified (n=1/18, 5.6%).

FGFR2 alterations were noted in three resistant biopsies (all with intrinsic resistance) (n=3/41, 7.3%), while activating mutations or amplification of ERBB2 was noted in five resistant biopsies (n=5/41, 12.2%). A single sensitive biopsy with an ERBB2 alteration was also identified (n=1/18, 5.6%).

With respect to ER signaling, three resistant biopsy samples exposed to CDK4/6i and an anti-estrogen demonstrated loss of ER expression via IHC (n=3/41, 7.3%); there were no patients with ER loss among the sensitive tumor samples (Figure 1B; Supplemental Table 5). These results support pre-clinical work suggesting CDK4/6i was predominantly effective in HR+ luminal cell lines while HR-basal cell lines demonstrated frequent intrinsic resistance.^11^

Enrichment in *ESR1* mutations was appreciated amongst resistant tumors (n=14/41, 34.1%; Supplemental Table 4) compared to sensitive tumors (n=3/18, 16.7%). *ESR1* mutations among sensitive tumors occurred exclusively in patients receiving fulvestrant and were not found in patients who achieved clinical benefit with CDK4/6i and an aromatase inhibitor, as would be expected (Supplementary Figure 1).^30^ These results support the notion that *ESR1* mutations are frequently acquired during the development of endocrine resistance, while also suggesting that they are not sufficient to drive simultaneous resistance to CDK4/6i.

Notably, mutational events in *PIK3CA* occurred frequently in both sensitive (n=8/18, 44.4%) and resistant (n=18/41, 43.9%) specimens, suggesting that *PI3KCA* is unlikely to be a marker of resistance. Copy number gains in *FGFR1* were also noted amongst both sensitive (n=4/18, 22.2%) and resistant biopsies (n=4/41, 9.8%).

Systematic differences in the relative proportion of these alterations were not apparent when comparing the intrinsic and acquired resistance subgroups, although the power of this analysis is limited by sample size (Figure 1D, Supplemental Table 4).

### Evolutionary dynamics in acquired CDK4/6i resistance

Matched pre- and post-treatment samples were available from seven patients who experienced acquired resistance to CDK4/6i. We compared the WES from the paired pre-treatment and post-treatment samples and performed an evolutionary analysis to evaluate clonal structure and dynamics, including changes in mutations and copy number. We established the evolutionary classification of each mutation to distinguish events that were acquired or enriched in clones that are dominant in the post-progression tumor, as compared with the pre-treatment counterpart (Figure 2, Supplemental Table 6).

**Figure 2.**
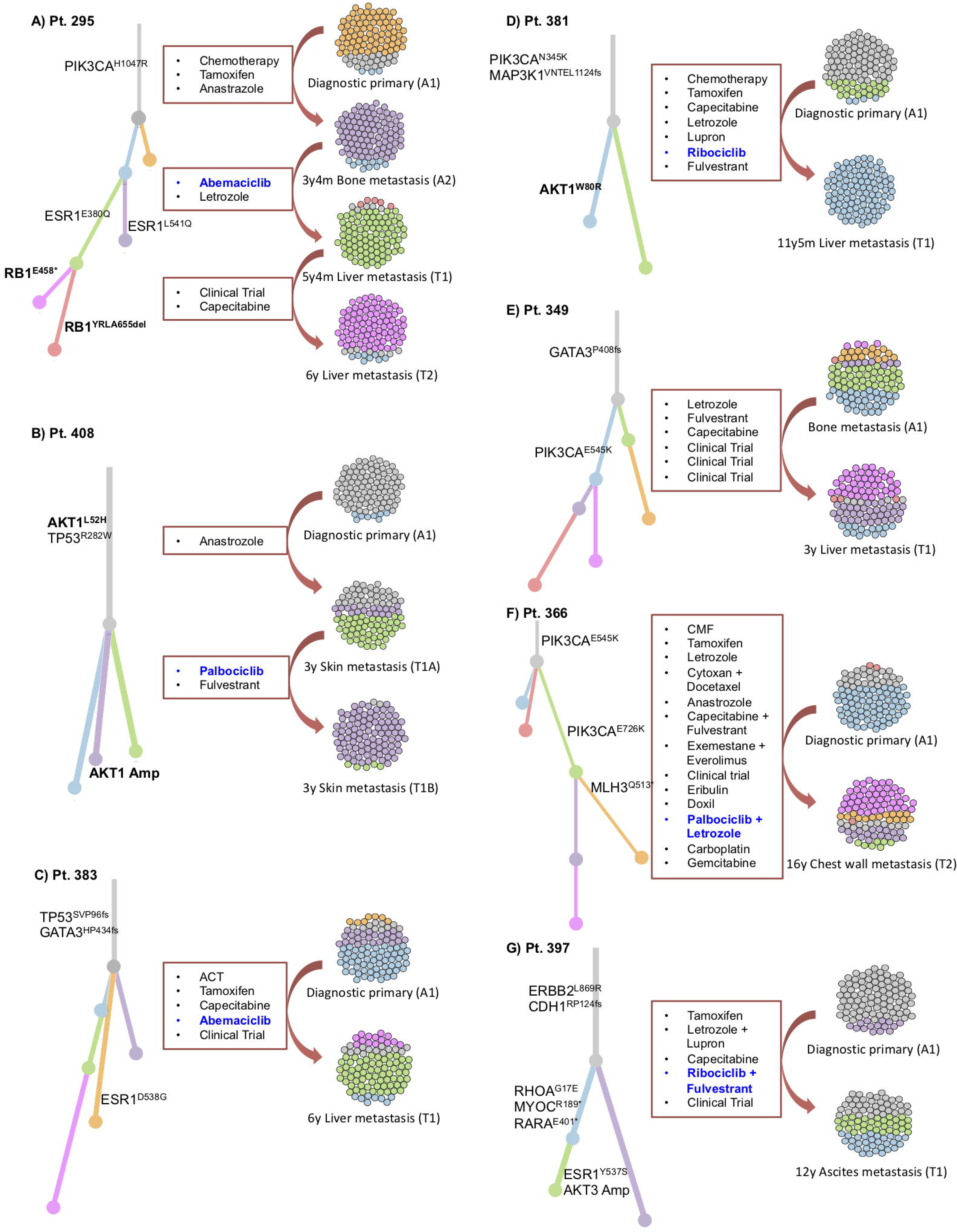
Acquired resistance to CDK4/6i in patients with pre-treatment and post-progression biopsies demonstrates convergent evolution of biallelic RB1 disruption and evolved AKT1 activation. Phylogenetic analysis depicting the evolutionary history for seven patients with acquired alterations, with clonal evolutionary dynamics demonstrating: (a) acquired polyclonal ESR1 mutations after aromatase inhibition, followed by convergent evolution of RB1 inactivation, with different RB1-inactivating mutations acquired in two parallel sibling clones; (b) Acquired AKT1 amplification; (c) No notable candidate for acquired mechanism of resistance (MOR); (d) Acquired AKT1 (W80R) mutation; (e) No notable candidate for acquired MOR; (f) Acquired inactivation of DNA Mismatch Repair Protein (MLH3); and (g) Acquired activating ESR1 mutation (Y537S) and amplification in AKT3.

Potential drivers of resistance that are observed in evolutionary acquired clones included a biallelic *RB1* disruption (Figure 2A), an *AKT1* amplification (Figure 2B), an *AKT1* activating mutation (Figure 2D), and an *ESR1* activating mutation (Figure 2G).

In the patient with biallelic *RB1* disruption and an available matched pair for exome analysis, the pre-treatment specimen demonstrated a single copy deletion in *RB1*. Two separate post-progression biopsy samples demonstrated unique alterations in the second copy of *RB1,* suggesting convergent evolution toward a common mechanism of resistance within the same tumor ecosystem (Figure 2A).

Genomic diversity was also observed in various mechanisms of AKT activation. In two patients with matched pre/post-treatment exome pairs, we observed acquisition of a pathogenic *AKT1* point mutation (*W80R*)^31–33^ (Figure 2D) and acquisition of an *AKT1* copy-number amplification (Figure 2B). Taken together, these cases suggest that cancer clones with activated AKT by either pathogenic mutation or high copy-number can confer selective advantage under CDK4/6i treatment.

In four of these pairs, the mechanism of acquired resistance remains unclear. We did not identify any instances of acquired AURKA overexpression, RAS activation, or CCNE2 amplification, though the analysis was limited by number of available matched pairs.

### Clinical case histories of patients with CDK4/6 inhibitor resistance

Figure 3 illustrates the clinical details of selected patients with intrinsic and acquired resistance to CDK4/6i and putative driver alterations. These include four instances of biallelic RB1 disruption (Figure 3A), three patients with AKT1 activation (Figure 3B), three with RAS activation (Figure 3C), and three with high CCNE2 amplification (Figure 3D).

**Figure 3.**
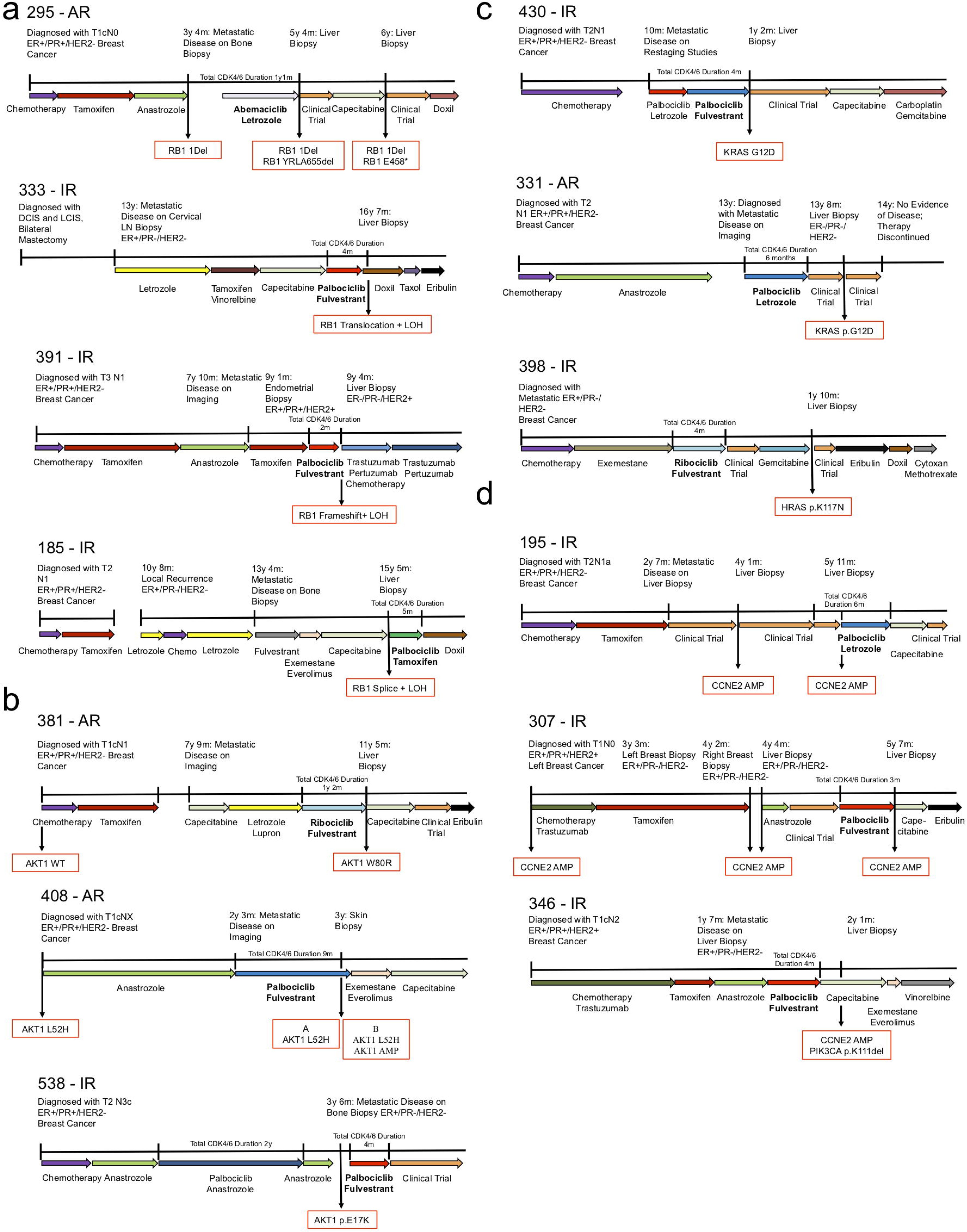
Clinical vignettes for candidate resistance drivers in representative patients (RB1, AKT1, RAS, and CCNE2). Clinical vignettes including treatment sequence, timing of metastatic progression, and available biopsies with key genomic findings are provided for the following - (a) four patients with biallelic alterations in RB1, including a patient with multiple biopsies and convergent evolution toward RB1 disruption (top, phylogenetic analysis for this patient is provided in Figure 2A). (b) Three patients with acquired alterations in AKT1 following progression on CDK4/6i. In the first (top), a new mutation in AKT1 W80R was identified. In the second (middle), a baseline alteration (AKT1 L52H) was identified at the time of diagnosis; at the time of progression on CDK4/6i, two biopsies were obtained – both demonstrating the baseline AKT1 L52H mutation, one also demonstrating an acquired amplification of the wild-type AKT1 protein (phylogenetic analyses for these patients are provided in Figure 2B and D). (c) Three patients with resistance to CDK4/6i and RAS-family alterations (including two instances of KRAS G12D and one instance of HRAS mutation). (d) Three patients with intrinsic resistance to CDK4/6i and amplification events in CCNE2.

Supplemental Figure 3 illustrates the three sensitive biopsy counter-examples: a single instance of AKT1 activation (Supplemental Figure 3A), a patient with low-level CCNE2 amplification (Supplemental Figure 3B), and a single ERBB2 alteration, all with clinical benefit on CDK4/6i (Supplemental Figure 3C).

Given the prominent (or exclusive) enrichment of *RB1* disruption, *AKT1* activation, RAS mutation, *AURKA* amplification, and *CCNE2* amplification within samples demonstrating resistance to CDK4/6i, we opted to pursue additional molecular validation of these targets. Prior work from our group and others implicating *FGFR* pathway and *ERBB2* activation in CDK4/6i resistance have been reported elsewhere.^24–26^

### Candidate alterations provoke resistance to CDK4/6i and anti-estrogens *in **vitro***

T47D and MCF7 HR+/HER2-breast cancer cells were utilized to explore whether these five genetic alterations confer resistance to CDK4/6i *in vitro*. *AKT1*, *KRAS G12D*, *AURKA,* and *CCNE2* were overexpressed via lentiviral transduction; *RB1* was inactivated via CRISPR-mediated knockout (Figure 4A; Supplemental Figure 4A). The impact of these alterations on susceptibility to CDK4/6 inhibitors was examined. Consistent with sequencing results, all alterations were sufficient to cause resistance to either palbociclib or abemaciclb in T47D cells (Figure 4B-F). Corresponding IC50 estimates for each dose-response curve are provided (Supplemental Table 7). Similar results were obtained in MCF7 cells (Supplemental Figure 3), though AURKA did not provoke resistance to CDK4/6i in this cell line, suggesting that context dependence may explain differences between cell lines, as with biopsies.

**Figure 4.**
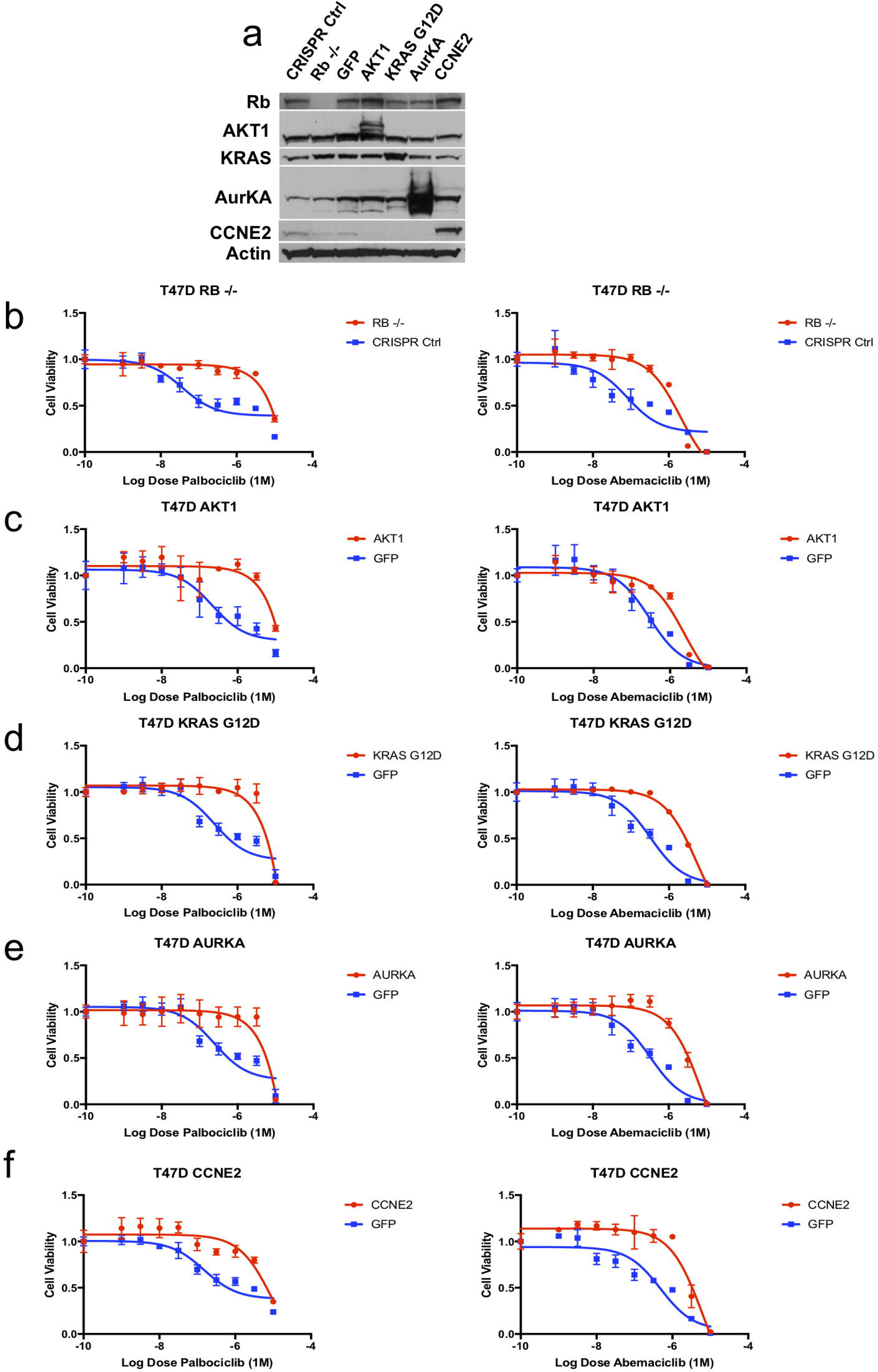
Candidate genomic alterations provoke CDK4/6i resistance *in vitro*. (a) T47D cells were modified via CRISPR-mediated downregulation (RB1) or lentiviral overexpression (AKT1, KRAS G12D, AURKA, CCNE2) to interrogate potential resistance mediators identified in patient biopsy samples. Western blotting with the indicated antibodies is included. (b-f) Modified T47D cells were exposed to escalating doses of CDK4/6i (palbociclib – left, abemaciclib – right) and viability was estimated via cell-titer-glo (CTG) assay. Control (CRISPR non-targeting guide or GFP) cells are plotted along with the resistance driver of interest (RB1 – b, AKT1 – c, KRAS G12D – d, AURKA – e, CCNE2 – f). Parental and variant cell lines are normalized to vehicle control and viability is plotted as a function of increasing CDK4/6i (graphed as triplicate average +/-standard deviation). All variants provoke CDK4/6i resistance (to both palbociclib and abemaciclib) *in vitro* in T47D cells. Corresponding IC50 values are included in Supplemental Table 7.

Given that most patients in the clinic are treated with a combination of CDK4/6i and an anti-estrogen, we also explored sensitivity to fulvestrant (Supplemental Figure 5). Cells lacking RB1 were only minimally resistant to fulvestrant monotherapy in both T47D and MCF7. Both AKT1 and CCNE2 overexpression conveyed resistance to fulvestrant in T47D and MCF7. Both KRAS G12D and AURKA overexpression provoked significant resistance to fulvestrant in T47D cells and but not in MCF7 cells.

Taken together, these results underscore the biological complexity related to the emergence of clinical resistance to these drug combinations both *in vitro* and in patients. They suggest that the resistance mechanisms identified in patient samples may provoke differential resistance to the CDK4/6- and estrogen-based components of the treatment regimen, and that these effects may depend upon additional cell-specific features.

### Resistance mediators arise independently during culture to resistance and define new dependencies *in vitro*

Given the results identified via exogenous manipulation of the mediators described above, we sought to explore resistance to CDK4/6i via orthogonal platforms in the laboratory. The HR+ cell lines T47D, MCF7, and MDA-MB-361 were cultured to resistance in the presence of increasing doses of palbociclib or abemaciclib. To examine whether the putative drivers identified in patients were also responsible for resistance under selection *in vitro*, we characterized the resistant derivatives for levels of retinoblastoma protein, aurora kinase, cyclin E2 and for activated effectors of KRAS or AKT1 (Figure 5A).

**Figure 5.**
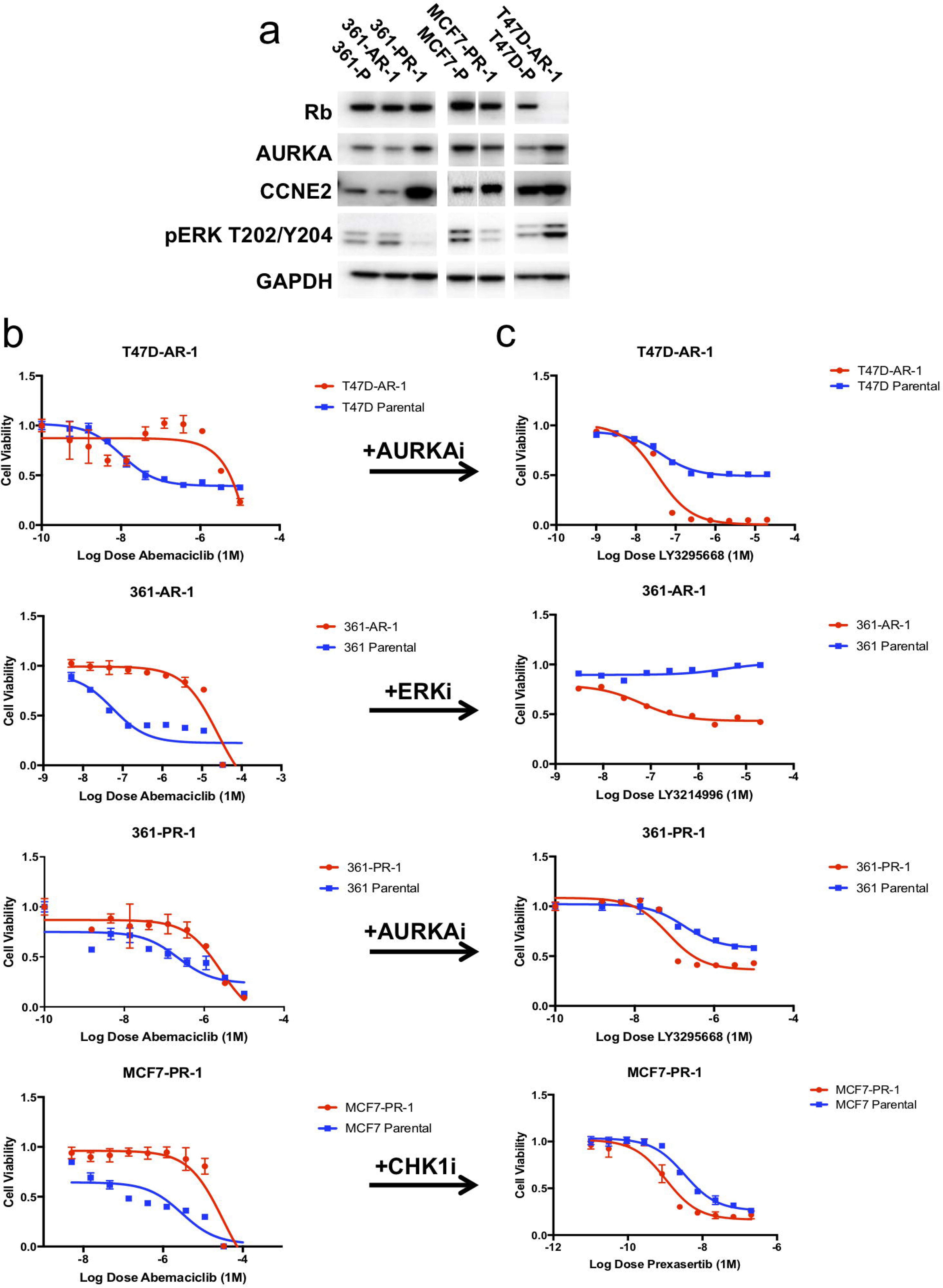
Candidate mutations emerge in cell lines cultured under CDK4/6i selective pressure and define new therapeutic dependencies *in vitro*. (a) Breast cancer cell lines (T47D, MCF7, MDA-MB-361) were cultured long-term to resistance in the presence of CDK4/6i (palbociclib, abemaciclib). The resulting cell lines which emerged were subjected to western blotting for putative mediators of drug resistance (RB1, AKT1, KRAS/ERK, AURKA, and CCNE2). (b-c) T47D cells cultured to resistance in the presence of abemaciclib demonstrated low levels of RB1 expression (T47D-AR1) and increased sensitivity to the AURKA inhibitor LY3295668. MDA-MB-361 cells cultured to resistance in the presence of abemaciclib demonstrated high levels of ERK activation (361-AR1) and increased sensitivity to the ERK inhibitor LY3214996. MDA-MB-361 cells cultured to resistance in the presence of palbociclib demonstrated high levels of AURKA (361-PR1) and increased sensitivity to the AURKA inhibitor LY3295668. MCF7 cells cultures to resistance in the presence of palbociclib demonstrated increased levels of CCNE2 (MCF7-PR1) and increased sensitivity to the CHEK1 inhibitor prexasertib.

Many of the putative resistance drivers identified via patient sequencing emerged spontaneously under selective pressure *in vitro*. 361-AR-1 (a derivative of MDA-MB-361 cells cultured to resistance in abemaciclib) was found to have an oncogenic *KRAS G12V* mutation (data not shown) and demonstrated increased ERK activation (Figure 5A). Proteomic analysis showed activation of multiple MAPK pathway components, including ERK, MEK and RSK (Supplemental Figure 6). T47D-AR-1 (a derivative of T47D cells cultured to resistance in abemaciclib) demonstrated decreased RB1 along with increased AURKA and pERK (Figure 5A). 361-PR-1 (a derivative of MDA-MB-361 cells cultured to resistance in palbociclib) demonstrated increased AURKA and CCNE2 protein levels (Figure 5A). Finally, MCF7-PR-1 (a derivative of MCF7 cells cultured to resistance in palbociclib) demonstrated increased expression of CCNE2 (Figure 5A). All derivative cell lines were confirmed to be resistant to abemaciclib compared with their parental counterparts (Figure 5B).

Therapeutic approaches are suggested by alterations identified in patient tumor specimens and cell lines cultured to resistance (Figure 5C). 361-AR-1 cells demonstrated increased KRAS/ERK activity and enhanced sensitivity to LY3214996, a selective ERK inhibitor. Both AURKA-amplified and RB1-low cells (T47D-AR-1 and 361-PR-1) were highly sensitive to LY3295668, a novel and selective AURKA inhibitor that has previously been reported to show synthetic lethality with RB1 loss.^34^ Finally, cancers with high cyclin E and CDK2 activation have been reported to be dependent on CHEK1.^35^ CCNE2-amplified cells (MCF7-PR-1) were highly sensitive to prexasertib, a CHEK1 inhibitor. Corresponding IC50 values for CDK4/6i and targeted agent treatment for these cell lines are included in Supplemental Table 8.

When compared to tumor sequencing results from patients with progression on CDK4/6i, the spontaneous emergence of corresponding alterations *in vitro* lends further support to the roles RB1 loss, RAS activation, CCNE2 overexpression, and AURKA overexpression may play in mediating resistance. That these alterations arose in parallel in different cancer cell lines (akin to different patients) also supports the earlier observation that cellular context may dictate which alterations arise under selective pressure via CDK4/6i. These results suggest that, in the presence of specific driver alterations in resistant tumor cells, unique dependencies may emerge which could inform novel therapeutic strategies.

### AURKA inhibition resulted in prolonged clinical benefit in a patient with HR+/HER2-, RB1+ MBC following progression on CDK4/6i-based therapy

LY3295668, the same AURKA specific inhibitor utilized *in vitro* to demonstrate a new dependence on AURKA in MDA-MB-361 and T47D cells cultured to resistance in CDK4/6i (Figure 5B, C), has entered early-stage clinical trials (NCT03092934).

As a proof-of-concept example, we provide the case history of a patient with locally advanced HR+/HER2-breast cancer treated on the trial. She had chemotherapy and adjuvant tamoxifen prior to metastatic recurrence; at that time, she was treated with first-line palbociclib and letrozole (Figure 6A). After prolonged clinical benefit on this regimen (>3 years), she progressed and enrolled on study with LY3295668. Her first restaging studies demonstrated disease stability, which persisted for approximately 11 months (Figure 6A, top). Immunohistochemical staining of her pre-treatment liver biopsy following progression on CDK4/6i demonstrated high levels of the proliferative marker Ki67 and high RB1 protein expression (Figure 6A, bottom), suggesting the mechanism of sensitivity to AURKA inhibition was not due to Rb loss. Sufficient additional biopsy material was not available for further sequencing or IHC-based analysis at the time of this writing. Our results lead us to speculate that sensitivity to AURKA inhibition in this patient could be due to alternative resistance mechanisms, such as AURKA amplification.

**Figure 6.**
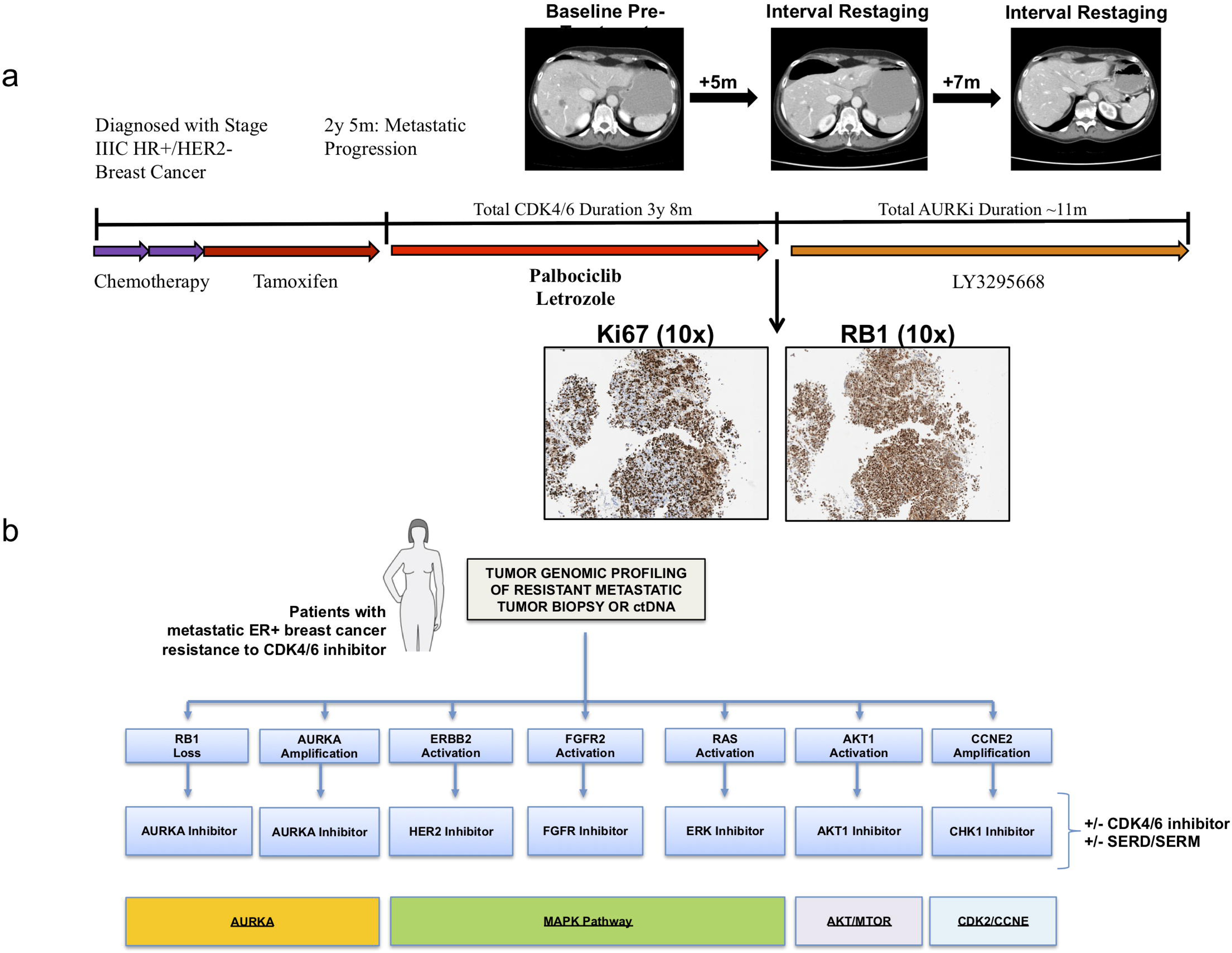
A novel aurora kinase A inhibitor demonstrates therapeutic efficacy in a patient with metastatic HR+ breast cancer after progression on CDK4/6i. (a) A patient with locally advanced HR+/HER2-breast cancer developed metastatic recurrence on adjuvant tamoxifen. She received CDK4/6i and letrozole in the first line setting with prolonged clinical benefit (>3 years). At progression, she was placed on trial with the AURKA inhibitor LY3295668; she subsequently experienced prolonged disease control ∼11 months. Baseline staging studies following progression on CDK4/6i in the patient described are included (top); she had osseous metastatic disease and visceral disease limited to the foci noted in the liver. Two interval restaging studies (top) demonstrate disease stability/mild response. Liver biopsy obtained at the time of progression on CDK4/6i and prior to LY3295668 demonstrated high Ki67 and high RB1 protein expression via immunohistochemistry (IHC, 10x) (bottom). (b) Schematic diagram demonstrating the potential utility of next-generation sequencing following progression on CDK4/6i; actionable alterations in RB1, ERBB2, FGFR2, AKT1, RAS, AURKA, and CCNE2 could dictate informed selection of targeted therapies as indicated.

## Discussion

CDK4/6 inhibitors, in combination with an anti-estrogen, have emerged as the standard of care for HR+/HER2-MBC. Despite widespread use, we have limited understanding of the mechanisms governing resistance and deciphering that landscape constitutes a critically important unmet need. To our knowledge, we provide the first analysis based upon whole exome sequencing of sensitive and resistant tumor tissues in a diverse cohort of patients who received CDK4/6i. This effort confirmed previous reports implicating rare events in RB1 while also revealing novel mediators of resistance including AKT1, RAS family oncogenes, AURKA, CCNE2, and ER loss. Prior work from our group and others identified mutational events in ERBB2^25^ and the FGFR pathway^24, 26, 36^ in driving resistance. *In vitro* experiments confirm that AKT1, KRAS G12D, AURKA, and CCNE2 confer resistance to CDK4/6i. RB1 downregulation, RAS/ERK activation, AURKA overexpression, and/or CCNE2 overexpression emerged spontaneously with prolonged CDK4/6i exposure, lending further support to their putative role as resistance effectors. These alterations correspond with the emergence of novel dependencies *in vitro*, providing therapeutic rationale for new targeted strategies in the clinic (Figure 6B). Finally, we provide an example of sustained clinical benefit with a novel AURKA inhibitor in a patient with HR+/HER2-MBC following progression on CDK4/6i.

Despite its central role downstream of CDK4/6, alterations in RB1 were observed only in a minority of patients who developed resistance to CDK4/6i. Anecdotal evidence of acquired alterations in RB1 at the time of progression was provided via ctDNA sequencing in three patients with exposure to CDK4/6i.^13^ ctDNA analysis from 195 patients treated on the PALOMA-3 study with fulvestrant and palbociclib also demonstrated rare RB1 alterations (∼5%), uniquely enriched in the palbociclib-containing arm.^14^ Relatively frequent driver alterations in *PIK3CA* and *ESR1* were also identified, though occurred in both treatment groups on PALOMA-3. These results were consistent with a recent study in which pre-treatment biopsies were subjected to targeted sequencing; alterations in RB1 were again rare (∼3%) and were associated with significantly impaired PFS on CDK4/6i.^22^ Our data supports the notion that RB1 alterations occur in a minority of CDK4/6i-resistant patients (4/41, ∼9.8%) and we provide new insight into diverse mechanisms of biallelic disruption. In a single patient with multiple pre- and post-treatment biopsies, two distinct mechanisms of biallelic inactivation were identified in separate post-progression biopsies, demonstrating convergent evolution under selective pressure for tumors with single copy loss *in vivo*. These findings were supported by culture to resistance experiments, in which multiple cell lines downregulated RB1 expression under selective pressure. While the rate of genomic RB1 disruption in tumor samples appears to be low following progression, additional non-genomic events may be missed by targeted or exome-based sequencing (such as methylation, mutations in regulatory regions, or post-translational modification). These possibilities warrant additional study.

Prior efforts suggested that common alterations in *CCND1*, *PIK3CA*, and *ESR1* did not impact PFS on CDK4/6i. We did not find an association between *CCND1*, *PIK3CA*, or *ESR1* alterations and CDK4/6i resistance in tumor specimens. Of note, alterations in *TP53* were enriched in CDK4/6i resistant biopsies. Mutant *TP53* is not sufficient to promote resistance to CDK4/6i *in vitro* as MCF7 (*TP53* wild-type) and T47D (*TP53* mutant) are both sensitive at baseline. Enrichment of *TP53* mutation in resistant specimens may result from heavier pre-treatment (including chemotherapies), may be permissive for the development of other resistance-promoting alterations, or may cooperate with secondary alterations to drive CDK4/6i resistance *in vivo*. The role of TP53 in CDK4/6i resistance remains an active area of research in the laboratory.

Several lines of evidence suggest CDK6 as a potential mechanism of resistance to CDK4/6 inhibitors.^19^ While clinical studies have not identified any examples of CDK6 alterations in resistant samples, a recent study that performed targeted sequencing in 348 tumor specimens obtained prior to treatment with CDK4/6i demonstrated that loss of function mutations in the *FAT1* tumor suppressor resulted in resistance to CDK4/6i. Interestingly, *FAT1* was shown to result in upregulation of CDK6 expression via the Hippo pathway *in vitro.*^22^. Finally, recent work from our institution demonstrated that micro-RNAs modulate CDK6 expression via the TGF-B pathway to alter sensitivity to CDK4/6i *in vitro.*^23^ Increased expression of the implicated miRNA (432-5p) correlated with resistance in a subset of the breast cancer patients exposed to CDK4/6i from the cohort analyzed here.^23^ In our study, we did not find examples of activating events in CDK6, nor did we identify *FAT1* alterations amongst resistant samples. Deletion and truncation mutations in *FAT1* appear to be extremely rare (reported in 6 of 348 patients in *Li et al*).^22^ Given their very low frequency and our sample size (n=58 patients), our study was likely not sufficiently powered to identify this rare event.

Unlike ctDNA-based targeted sequencing reported from the PALOMA-3 study, the cohort analyzed here represents, to our knowledge, the first analysis based upon whole exome sequencing from clinically annotated biopsies reflecting a diverse group of patients with exposure to multiple CDK4/6i-based regimens. In addition to expected alterations in RB1, we identified a heterogeneous landscape of resistance, in which a variety of rare driver events span a diverse spectrum of potential mediators. We confirm enrichment of activating mutations in *ERBB2* and amplification events in *FGFR2* in resistant patients, and both pathways provoke resistance to anti-estrogens and CDK4/6i *in vitro.*^24–26, 36^ We present, to our knowledge, the first evidence implicating AKT1, RAS, and AURKA in mediating resistance to CDK4/6i in patients. Targeted sequencing of ctDNA via samples from PALOMA-3 also identified rare events in *ERBB2*, *AKT1*, *KRAS*, and *FGFR2* which were both acquired and maintained at progression, however this analysis was limited by lack of insight into the clinical response phenotype of these samples.^14^ We would hypothesize that “maintained” alterations identified in the context of that study represent instances of early- or intrinsic resistance while “acquired” alterations are more likely to arise in patients with transient response or clinical benefit from CDK4/6i. CCNE2 and AURKA did not emerge as potential resistance mediators in that study, likely due to lack of insight into copy number alterations as a result of the sequencing methodology.

More recent correlative analyses from PALOMA-3 suggested that CCNE1 expression is associated with inferior outcome for patients receiving palbociclib.^21^ While we did not see examples of CCNE1 amplification in this cohort, we do provide, to our knowledge, the first evidence that CCNE2 amplification is also associated with the resistant phenotype. Of note, given its proximity to the centromere, copy number analysis of CCNE1 via WES is technically challenging and this may have resulted in under-estimation of amplification events in this gene.

While all of these mediators provoked resistance to CDK4/6i *in vitro*, in specific instances there were cell-line-dependent differences in their ability to circumvent CDK4/6i. This notion of context-specificity is supported by several isolated counter-examples in patients, in which putative resistance mediators were found to occur in individual patients who derived at least transient clinical benefit from CDK4/6i. These findings are also consistent with the spontaneous emergence of distinct resistance mediators in specific cell lines – for example, RAS/ERK-activated and AURKA-amplified cells emerged in MDA-MB-361 but not in MCF7, and exogenous overexpression of AURKA could not provoke resistance in MCF7. The situation is further complicated by variation in anti-estrogen resistance *in vitro*. As an example, AKT1 overexpression may be sufficient to provoke resistance to both CDK4/6i and fulvestrant, while alterations in RB1 may require a second cooperative event to overcome the anti-estrogen component of the regimen (such as *ESR1* alteration). These nuances underscore the complexity of modeling resistance to therapeutic combinations *in vitro* and highlight the need for additional studies to explore context-specific factors, which might dictate the emergence of resistance with a potential driver of interest.

The majority of alterations identified in our clinical cohort, and confirmed *in vitro*, are amenable to therapeutic intervention via emerging agents (Figure 6B). These results suggest that a non-selective regimen is unlikely to yield reliable clinical benefit, while a precision-based approach, informed by the underlying genomic findings at progression, could guide selection of therapy in CDK4/6i-resistant patients. RAS-activated cells that emerged under selective pressure with CDK4/6i were highly sensitive to LY3214996, a selective ERK inhibitor. The CHEK1 kinase plays well-established roles in regulating cell cycle progression in the setting of DNA damage.^37^ Cancer cells with replication stress caused by activated CDK2 appear to be particularly sensitive to Chk inhibitors^38^ and *CCNE1* amplification has been linked to CHEK1 dependence.^35^ HR+ cells expressing high levels of CCNE2 demonstrated enhanced sensitivity to prexasertib, a CHEK1 inhibitor that has been well tolerated in human patients with early evidence suggesting clinical efficacy in a phase I study.^39^

The aurora kinases regulate organization of the mitotic spindle and cell cycle progression.^40^ AURKA overexpression in breast cancer has been associated with an ER-low/basal phenotype.^41^ AURKA was previously implicated in mediating endocrine resistance via SMAD-dependent downregulation of ER-alpha expression.^42^ Alisertib, an oral AURKA inhibitor, was well tolerated in HR+ MBC patients when combined with fulvestrant, and anti-tumor activity was appreciated in a phase I trial.^43^ A randomized phase II study of this combination has completed accrual (NCT02860000). We demonstrate that HR+ cells cultured to resistance in CDK4/6i can demonstrate downregulation of RB1 or increased expression of AURKA, both of which are associated with increased sensitivity to LY3295668, a novel selective AURKA inhibitor. In screens to identify synthetic lethal interactions with an RB1 mutation in lung and other cancers, the aurora kinases emerged as key targets, and LY3295668 provoked tumor regression in xenograft models of RB1-null small cell lung cancer.^34, 44^ We provide the first evidence supporting AURKA as a mediator of resistance to CDK4/6i *in vitro* and in tumor samples. Furthermore, in a patient with HR+ MBC who progressed after a prolonged course of CDK4/6i-based therapy (analogous to our translational culture-to-resistance experiment *in vitro*), subsequent treatment on a phase I trial with LY3295668 was well tolerated and prompted prolonged clinical benefit. This patient had high RB1 protein expression at the time of therapy initiation, suggesting that her response was not governed by RB1 loss. Based upon these translational insights, a phase I study exploring the utility of LY3295668 in patients with HR+ MBC following progression on CDK4/6i was recently initiated (NCT03955939).

Although one can consider targeting each individual resistance mechanism directly, it may also be possible to target a smaller number of resistance “nodes” or pathways upon which multiple resistance effectors converge. We previously showed that *ERBB2* mutations and alterations in FGFR1/FGFR2 activate the MAPK pathway in resistant HR+ MBC,^24, 25^ and MAPK pathway inhibition was able to overcome this resistance. *RAS* mutations also activate the MAPK pathway. The fact that multiple mechanisms of resistance to CDK4/6i activate the MAPK pathway suggests that this may be an important node of resistance in HR+ MBC – and that combining endocrine therapy and CDK4/6i with agents that target MAPK such as MEK inhibitors, ERK inhibitors, and/or SHP2 inhibitors, may be a unifying strategy to overcome or prevent resistance resulting from multiple genetic aberrations. Similarly, both RB loss and AURKA amplification are targetable with AURKA inhibitors. Taken together, it may be possible to address all seven of these mechanisms (which account for at least 80% of the resistant biopsies in this study) by targeting four nodes/pathways: AURKA, MAPK, AKT/MTOR, and CCNE/CDK2 (Figure 6B).

We have identified multiple novel effectors of resistance to CDK4/6i in HR + breast cancer, providing rationale to guide the development of a wide range of precision-based clinical trials, in which patients with specific genomic or molecular alterations are treated with novel therapeutic combinations designed to circumvent or overcome resistance.

## Methods

### Patients and Tumor Samples

Prior to any study procedures, all patients provided written informed consent for research biopsies and whole exome sequencing of tumor and normal DNA, as approved by the Dana-Farber/Harvard Cancer Center Institutional Review Board (DF/HCC Protocol 05-246). Metastatic core biopsies were obtained from patients and samples were immediately snap frozen in OCT and stored in -80°C. Archival FFPE blocks of primary tumor samples were also obtained. A blood sample was obtained during the course of treatment, and whole blood was stored at -80°C until DNA extraction from peripheral blood mononuclear cells (for germline DNA) was performed. In a few instances, cell free DNA was obtained from plasma for circulating tumor DNA analysis, as previously described.^49^

### Clinical Annotation and Biopsy Phenotypes

Patient charts were reviewed to determine the sequence of treatments received in the neoadjuvant, adjuvant, and metastatic setting as well as the temporal relationship between available biopsy samples and CDK4/6i exposure. Radiographic parameters were assigned via review of the imaging study interpretations available in the patient record during the CDK4/6i treatment course – tumors were defined as “responding” if any degree of tumor shrinkage was reported by the evaluating radiologist, “stable” if there was felt to be no meaningful change, “progressing” if lesions were increasing in size, or “mixed” if comment was made denoting simultaneous shrinkage and growth in discordant lesions. Tumors with a mixed response were excluded from analysis as a reliable phenotype could not be assigned. The “best radiographic response” (BRR) was then assigned as either “response” (R), “stable disease” (S), or “progression” (P) based upon the best radiographic parameter noted during the CDK4/6i treatment course.

Sensitive biopsies were defined as baseline samples obtained within 120 days prior to, or up to a maximum of 31 days after, CDK4/6i treatment initiation in a patient with subsequent clinical benefit (radiographic response or stable disease >6 months). Biopsies reflecting acquired resistance were obtained from patients who had experienced clinical benefit with CDK4/6i and had an available biopsy specimen either within 31 days of progression or at any time thereafter. Biopsies reflecting intrinsic resistance were obtained within 120 days prior to CDK4/6i initiation in patients without evidence of clinical benefit (defined as progression on the first interval restaging study or stable disease <6 months).

### Whole Exome Sequencing

DNA was extracted from primary tumors, metastatic tumors, and peripheral blood mononuclear cells (for germline DNA) from all patients and whole exome sequencing was performed, as detailed below. In several instances, cell free DNA was obtained from plasma for circulating tumor DNA analysis, as previously described.^49^

#### DNA extraction

DNA extraction was performed as previously described.^50^ For whole blood, DNA is extracted using magnetic bead-based chemistry in conjunction with the Chemagic MSM I instrument manufactured by Perkin Elmer. Following red blood cell lysis, magnetic beads bind to the DNA and are removed from solution using electromagnetized rods. Several wash steps follow to eliminate cell debris and protein residue from DNA bound to the magnetic beads. DNA is then eluted in TE buffer. For frozen tumor tissue, DNA and RNA are extracted simultaneously from a single frozen tissue or cell pellet sample using the AllPrep DNA/RNA kit (Qiagen). For FFPE tumor tissues, DNA and RNA are extracted simultaneously using Qiagen’s AllPrep DNA/RNA FFPE kit. All DNA is quantified using Picogreen

#### Library Construction

DNA libraries for massively parallel sequencing were generated as previously described^50^ with the following modifications: the initial genomic DNA input into the shearing step was reduced from 3µg to 10-100ng in 50µL of solution. For adapter ligation, Illumina paired-end adapters were replaced with palindromic forked adapters (purchased from Integrated DNA Technologies) with unique dual indexed 8 base index molecular barcode sequences included in the adapter sequence to facilitate downstream pooling. With the exception of the palindromic forked adapters, all reagents used for end repair, A-base addition, adapter ligation, and library enrichment PCR were purchased from KAPA Biosciences in 96-reaction kits. In addition, during the post-enrichment solid phase reversible immobilization (SPRI) bead cleanup, elution volume was reduced to 30µL to maximize library concentration, and a vortexing step was added to maximize the amount of template eluted.

#### Solution-phase hybrid selection

After library construction, hybridization and capture were performed using the relevant components of Illumina’s Rapid Capture Exome Kit and following the manufacturer’s suggested protocol, with the following exceptions: first, all libraries within a library construction plate were pooled prior to hybridization. Second, the Midi plate from Illumina’s Rapid Capture Exome kit was replaced with a skirted PCR plate to facilitate automation. All hybridization and capture steps were automated on the Agilent Bravo liquid handling system.

#### Preparation of libraries for cluster amplification and sequencing

After post-capture enrichment, library pools were then quantified using quantitative PCR (KAPA Biosystems) with probes specific to the ends of the adapters; this assay was automated using Agilent’s Bravo liquid handling platform. Based on qPCR quantification, libraries were normalized and denatured using 0.1 N NaOH on the Hamilton Starlet.

#### Cluster amplification and sequencing

Cluster amplification of denatured templates was performed according to the manufacturer’s protocol (Illumina) using HiSeq 2500 Rapid Run v1/v2, HiSeq 2500 High Output v4 or HiSeq 4000 v1 cluster chemistry and HiSeq 2500 (Rapid or High Output) or HiSeq 4000 flowcells. Flowcells were sequenced on HiSeq 2500 using v1 (Rapid Run flowcells) or v4 (High Output flowcells) Sequencing-by-Synthesis chemistry or v1 Sequencing-by-Synthesis chemistry for HiSeq 4000 flowcells. The flowcells were then analyzed using RTA v.1.18.64 or later. Each pool of whole exome libraries was run on paired 76np runs, with a two 8 base index sequencing reads to identify molecular indices, across the number of lanes needed to meet coverage for all libraries in the pool.

#### Sequence data processing

Exome sequence data processing was performed using established analytical pipelines at the Broad Institute. A BAM file was produced with the Picard pipeline (see URLs) which aligns the tumor and normal sequences to the hg19 human genome build using Illumina sequencing reads. The BAM was uploaded into the Firehose pipeline (see URLs), which manages input and output files to be executed by GenePattern ^51^.

#### Sequencing quality control

Quality control modules within Firehose were applied to all sequencing data for comparison of the origin for tumor and normal genotypes and to assess fingerprinting concordance. Cross-contamination of samples was estimated using ContEst.^52^

### Somatic Alteration Assessment

MuTect^53^ was applied to identify somatic single-nucleotide variants. Indelocator (see URLs), Strelka^54^, and MuTect2 (see URLs) were applied to identify small insertions or deletions. A voting scheme with inferred indels requiring at least 2 out of 3 algorithms.

Artifacts introduced by DNA oxidation (so called OxoG) during sequencing were computationally removed using a filter-based method.^55^ In the analysis of primary tumors that are formalin-fixed, paraffin-embedded samples [FFPE] we further applied a filter to remove FFPE-related artifacts.^56^

Reads around mutated sites were realigned with Novoalign (see URLs) to filter out false positive that are due to regions of low reliability in the reads alignment. At the last step, we filtered mutations that are present in a comprehensive WES panel of 8,334 normal samples (using the Agilent technology for WES capture) aiming to filter either germline sites or recurrent artifactual sites. We further used a smaller WES panel of normal 355 normal samples that are based on Illumina technology for WES capture, and another panel of 140 normals sequenced within our cohort^27^ to further capture possible batch-specific artifacts. Annotation of identified variants was done using Oncotator.^57^

### Copy Number and Copy Ratio Analysis

To infer somatic copy number from WES, we used ReCapSeg (see URLs), calculating proportional coverage for each target region (i.e., reads in the target/total reads) followed by segment normalization using the median coverage in a panel of normal samples. The resulting copy ratios were segmented using the circular binary segmentation algorithm.^58^

To infer allele-specific copy ratios, we mapped all germline heterozygous sites in the germline normal sample using GATK Haplotype Caller^59^ and then evaluated the read counts at the germline heterozygous sites in order to assess the copy profile of each homologous chromosome. The allele-specific copy profiles were segmented to produce allele specific copy ratios.

### Gene deletions and Bi-allelic inactivation

For the inference of gene deletions and inactivations, as we aim to infer bi-allelic inactivations (BiDel or “HOMDEL”), we take into account various mutational events that may result in inactivation of both alleles. These mutational events include: (1) loss of heterozygosity (LOH), (2) SNV (while excluding the following variant classifications: “Silent”, “Intron”, “IGR”, “5’UTR”, “3’UTR”, “5’Flank”, “3’Flank”), (3) short indels, (4) long deletions and gene rearrangements inferred by SvABA,^60^ and (5) potentially pathogenic germline events in cancer genes (see description below).

Potentially pathogenic germline events: aiming to retain a subset of potentially pathogenic germline events there are several features which are accounted for including (1) ClinVar significant annotation among the following: Pathogenic. Likely pathogenic, Conflicting interpretations of pathogenicity, risk factor or (2) Variant Classification among the following: Splice_Site, Frame_Shift_Del, Frame_Shift_Ins, Nonsense_Mutation. In addition (3) Genome Aggregation Database (gnomAD)^61^ less than 0.05 (indicating it is a rare variant)

### Cancer Cell Fraction and Evolutionary Analysis

#### Analysis using ABSOLUTE

To properly compare SNVs and indels in paired metastatic and primary samples, we considered the union of all mutations called in either of the two samples. We evaluated the reference and alternate reads in each patient’s primary and metastatic tumors, including mutations that were not initially called in one of the samples. These mutations in matched samples were used as input for ABSOLUTE.^62^ The ABSOLUTE algorithm uses mutation-specific variant allele fractions (VAF) together with the computed purity, ploidy, and segment-specific allelic copy-ratio to compute cancer cell fractions (CCFs).

### Clonal structure and phylogenetic reconstruction of tumor evolution

The clonal structure observed in individuals with more than a single tumor sample was inferred with PyClone,^63^ using the Beta Binomial model and the copy number of each mutation inferred by ABSOLUTE with the parental copy number parameter.

Subsequently, the inferred clonal structure was used to trace the evolutionary history of the clones (phylogenic tree) using the ClonEvol,^64^ retaining only clones with at least four mutations and estimated cancer cellular fraction (cellular prevalence) higher than 1%.

### Evolutionary analysis of copy-number variation

#### Corrected quantification of copy number

gene amplifications are based on the purity corrected measure for the segment containing that gene, based on ABSOLUTE (rescaled_total_cn).^62^ To better measure segment-specific copy-number, we subtracted the genome ploidy for each sample to compute copy number above ploidy (CNAP). CNAP of at least 3 are considered as amplifications (“AMP”), CNAP above 1.5, but below 3 are considered low amplification (“GAIN”), and are not depicted in our mutational landscape (Figure 1). CNAP of at least 6 are considered high amplifications (“HighAMP”), and CNAP of at least 9 and no more than 100 genes^65^ is considered very high focal amplification (“FocalAMP”).

The evolutionary classification of amplifications accounts for the magnitude of the observed copy-number difference between the pre-treatment and the post-treatment samples. If the difference between the CNAP of the post-treatment and the CNAP of the pre-treatment is smaller than 50%, the amplification is defined as “Shared”. If the CNAP of the post-treatment is larger than the CNAP by more than 50% and the lower pre-treatment CNAP is not at “FocalAMP” level, the evolutionary classification is “Acquired”. If CNAP of the post-treatment is smaller by at least 50%, comparing to the pre-treatment sample and the lower post-treatment CNAP is not at “FocalAMP” level, the evolutionary classification is “Loss”. Otherwise, the evolutionary classification of amplifications is defined as “Indeterminate”.

### Cell Culture

HR+/HER2-human breast cancer cell lines T47D (HT-133) and MCF7 (HTB-22) were obtained from American Type Culture Collection (ATCC). T47D and MCF7 cells were cultured in RPMI 1640 medium (no phenol red; Gibco, 11835-030) and MEMα (nucleosides, no phenol red; Gibco, 41061029) respectively, both supplemented with 10% fetal bovine serum (Gemini bio-products, 100-106) and 1% penicillin-streptomycin-glutamine. HEK 293T/17 (CRL-11268) were obtained from ATCC and cultured in DMEM (high glucose, pyruvate; Gibco, 11995065), supplemented with 10% fetal bovine serum (Gemini bio-products, 100-106) and 1% penicillin-streptomycin-glutamine (Gibco, 10378016).

### Candidate driver plasmid and cell line production

AKT1 (BRDN0000464992), KRASG12D (BRDN0000553331), AURKA (TRCN0000492002), CCNE2 (ccsbBroadEn_11340), and GFP bacterial streaks were obtained from the Genetic Perturbation Platform, Broad Institute, MA. RB1 and CRISPR non-targeting guide cells were obtained as a gift Flora Luo and the Garraway laboratory. The CCNE2 construct was cloned into a pLX307 vector using the LR reaction kit (Life Technologies, 11791019). All construct plasmids were prepared using the Plasmid Plus Midi Kit (Qiagen, 12943). To generate lentivirus for each construct, 293T cells were transfected with Opti-MEM (Gibco, 31985-062), FuGENE HD (Promega, E2311), VSV-G envelope plasmid, and L8.91 packaging plasmid. After 72h of incubation, supernatant was filtered through a 0.45 μL filter (Corning, 431225) and lentivirus presence was tested using Lenti-X GoStix (TakaraBio, 631244). 500μL – 1mL of virus was added to a 60-mm dish containing T47D (or MCF7) cells and medium with 4μg/mL of polybrene (Millipore Sigma, TR-1003-G). After overnight incubation, cells were moved to a 100-mm dish and again incubated overnight. The medium was replaced and 0.5μg/mL of puromycin (Gibco, A1113803) were added to KRASG12D, AURKA, CCNE2, RB1 and CRISPR constructs, and 6-10μg/mL of blasticidin (Gibco, A1113903) were added to GFP and AKT1 constructs. Plates were compared to uninfected control plates, and after 2 days of selection, were plated for drug sensitivity assay and harvested for western blotting as described below.

### Kill Curves/Drug Sensitivity Assay

Cells were plated at a density of 1000 cells/well in RPMI and 1500 cells/well in MEMα, for T47D and MCF7, respectively, in 96 well plates (PerkinElmer, 6005181). The experiments were plated in triplicate, for ten doses of the drug of interest. Palbociclib doses ranging from 1 nM to 10 μM were prepared from a 10 mM stock solution in molecular biology grade water (Corning, 46-000-CI); abemaciclib doses ranging from 1 nM to 10 μM were prepared from a 10 mM stock solution in molecular biology grade water (Corning, 46-000-CI); fulvestrant doses ranging from 0.01 nM to 1 μM were prepared from a 20 mM stock solution in DMSO (Sigma-Aldrich, D2650). The next day, cells were treated with the range of doses of the drug of interest. Cells were re-treated three days later.

After treatment has been applied for eight days, the 96-well plates were brought out of the incubator and allowed to equilibrate to room temperature. The medium was replaced with 50 μL of fresh medium per well. 50 μL of CellTiter-Glo 2.0 (Promega, G9241) was added to each well, the plate was shaken at 200 rpm for min, and then allowed to equilibrate at room temperature for fifteen minutes as per the CellTiter-Glo 2.0 Assay Technical Manual. Average background luminesce reading was calculated from plate wells containing only medium, and was subtracted from all values. The values were then averaged for each triplicate and standard deviations were calculated. The data were normalized to the no-drug, vehicle control for each construct. The calculated averages and standard deviations were visualized on GraphPad Prism 7 using the log(inhibitor) vs. response (three parameters) preset protocol.

### Chemicals and antibodies

Chemicals utilized included palbociclib (Selleck Chemicals, S1116), abemaciclib (ApexBio, A1794), and fulvestrant (Sigma-Aldrich, I4409). Primary antibodies utilized included antibodies against β-Actin (Santa Cruz, sc-47778), Rb (Cell Signaling Technology, clone 4H1, 9309), Akt (CST, 9272), Ras (CST, clone D2C1, 8955), Aurora A (CST, clone D3E4Q, 14475), and Cyclin E2 (CST, 4132), in addition to the secondary antibodies goat anti-rabbit (Invitrogen, 32260) and goat anti-mouse (Invitrogen, A16090).

### Western blotting

A near-confluent T75 (∼7x10^6 cells) was spun down and the pellet kept at -20C. The pellet was then lysed in 1mL of lysis buffer consisting of RIPA buffer (Sigma-Aldrich, R0278), dithiothreitol (DTT, Invitrogen, 15508013), phenylmethane sulfonyl fluoride (PMSF, Sigma-Aldrich, P7626), and PhosStop (Sigma-Aldrich, 4906837001). Lysate was rotated at 15 r.p.m for 15 minutes at 4°C, then centrifuged at 14,000g for 15 minutes at 4°C, preserving the supernatant. Protein concentration was quantified via bicinchoninic acid assay (Pierce BCA Protein Assay Kit, Thermo Fisher Scientific, 23225) and Tecan i-control software pre-set BCA program. Samples were prepared using 40μg of protein, Bolt LDS Sample Buffer (Invitrogen, B0007), and DTT and heated to 95°C for 5 min. The samples were run on a Bolt 4-12% Bis-Tris Plus Gel (Invitrogen, NW04120BOX) in 1X Bolt MOPS SDS Running Buffer (Invitrogen, B000102) for 1hr at 130V. Protein was transferred to nitrocellulose membranes via the Trans-Blot Turbo Transfer System (Bio-Rad, 1704150) following the turbo mini preset protocol (1.3A 25V 7Min) two times. Membranes were blocked in 5% milk in Tris-buffered saline (Bio-Rad, 1706435) with 0.1% Tween-20 (Sigma-Aldrich, P9416) for one hour at room temperature. Membranes were incubated overnight at 4°C with primary antibodies that were diluted 1:1000 (with the exception of Rb, which was diluted 1:500) in 5% milk in TBS-T. After incubation, membranes were washed 3 times for 10min with 1X TBS-T and incubated with secondary antibody diluted 1:2000 in 5% milk in TBS-T for 1h at room temperature. Membranes were then washed times for 10min with 1X TBS-T. After washing, membranes were treated with Pierce ECL Plus Western Blotting Substrate (Thermo Fisher Scientific, 32132) for 5 minutes and exposed to autoradiography film (Denville, 1159M38).

For resistant/derivative cell lines: cells were washed with PBS and lysed in lysis buffer (1% triton X-100, 25mM Tris pH 7.5, 150mM NaCl, 1mM EDTA, Halt Protease/phosphatase inhibitor cocktail), and protein concentration was assessed by BCA protein assay (Pierce 23225). Equal amounts of protein were electrophoresed on 4-20% BioRad Tris Glycine Gels (BioRad 5671094) transferred to nitrocellulose (BioRad 1704159) and probed with primary antibodies. Antibodies were purchased from Cell Signaling Technology for Rb Total (9307), pRb S780 (3590), pRb S807/811 (8516), CCNE2 (4132), Akt S473 (4051), S6 total (2317), S6 S240/244 (4838), ERK total (3042), pERK T202/Y204 (4370, 4376) and R&D Systems AurA (AF3295). Digiwest® protein profiling of MDA-MB-361-AR was also conducted with NMI TT.

### Resistant cell line generation

The methods for generating resistant cell lines were described previously.^34^ Briefly, MDA-MB-361, T47D and MCF-7 ER+ breast cancer cell lines were used to derive variants with acquired resistance to abemaciclib or palbociclib. T47D (HTB-133), MCF-7 (HTB-22) and MDA-MB-361 (HTB-27) were purchased from The American Type Culture Collection (ATCC). Cell lines were cultured in RPMI-1640 medium (Gibco 22400-089) + 10% FBS (Hyclone SH30071.03), Eagles Essential Medium (Gibco 11090-081) + 10% FBS and Liebovitz L-15 Medium (Gibco 11215-064) + 20% FBS, respectively. Resistant cell lines were generated by chronic treatment with either abemaciclib or palbociclib alone or in combination with fulvestrant. Cell cultures were initiated in low doses of compound approximating the IC50 until cells grew to 80% confluence. Cells were then passaged and treated with incrementally higher doses. This process was repeated several times until cells were able to grow in the presence of drugs at clinically meaningful concentrations. Once resistant cell lines were established, the stability of resistance was assessed with a 21 day dosing holiday. Resistance remained stable in all cell lines except for T47D-AR and T47D-PR which became almost completely resensitized to the CDK4/6i after the 21 day drug-free period. All resistant derivatives resistant were found to be cross resistant to the CDk4/6i that was not used in the selection step. Short tandem repeat (STR) analysis was performed to verify the authenticity of the cell lines.

### Proliferation Assays

Cells were plated onto poly-D-lycine plates (Corning 354640) and treated in replicate with a dose curve of compounds of interest. Cells were allowed to grow for two doubling times and proliferation was measured by CellTiter-Glo® (Promega G7571) or CyQuant (Invitrogen C3511) per manufacturer’s protocol. Data analysis was carried out using Prism software.

### LY3295668 Phase 1/2 Clinical Trial

The patient vignette provided in this manuscript was shared from an ongoing phase 1/2 study. Please see protocol NCT03092934 at www.clinicaltrials.gov for details related to the study location, eligibility, and compound. This is an open-label, multicenter study of patients with locally advanced or metastatic solid tumors and disease progression after 1-L4 prior treatment regimens. The phase 1 portion of the protocol is designed to evaluate the primary objective of determining the maximum tolerated dose (MTD); secondary objectives included evaluation of tolerability and overall safety profile of LY3295668. The primary objective of the phase 2 study portion is to evaluate the objective response rate of tumors after treatment with LY3295668. Patients in the phase 2 study were required to have estrogen receptor and/or progesterone receptor positive, human epidermal growth factor receptor 2 (HER2) negative, breast cancer with prior exposure to and progression on on a hormone therapy and a CDK4/6 inhibitor.

## Acknowledgements

We thank Laura Dellostritto, Lori Marini, Nelly Oliver, Shreevidya Periyasamy, Janet Files, Sara Hoffman, and Colin Mackichan for assistance with patient sample collection and annotation. We are grateful to all the patients who volunteered for our tumor biopsy protocol and generously provided the tissue analyzed in this study.

## Supplementary Figure Legends

**Supplemental Figure 1. Subgroup genomic analysis of the CDK4/6i cohort based upon anti-estrogen exposure.**

Heatmaps demonstrating key genomic events (both copy number alteration and mutation) in a subset of genes for (a) patients with exposure to CDK4/6i and aromatase inhibitor (AI) and for (b) patients with exposure to CDK4/6i and fulvestrant. The gene set and clinical parameters are identical to those provided in Figure 1B.

**Supplemental Figure 2. Higher AURKA expression observed even in low-amplification tumors in TCGA**

Breast tumor from the TCGA dataset were stratified based on the genomic AURKA copy number (low amplification – left, no amplification – right; high amplification excluded) and plotted against AURKA RNA expression. Higher AURKA RNA expression was observed in low AURKA-amplification compared to non-amplified tumors in these TCGA samples.

**Supplemental Figure 3. Candidate resistance mutations in representative patients – key counterexamples.**

Biopsies demonstrating CDK4/6i sensitivity despite the presence of putative resistance drivers were identified and clinical vignettes were generated. (a) A patient with bone-only metastatic progression was placed on first-line CDK4/6i and letrozole. A canonical AKT1 E17K alteration was identified at the time of metastatic progression. This patient has had stable osseous metastatic disease on interval repeat imaging and remained on treatment at the time of data cutoff. A patient with de novo metastatic HR+/HER2-breast cancer was treated with tamoxifen and subsequently received palbociclib and letrozole. Prior to CDK4/6i exposure, which lasted for a duration exceeding one year, a baseline low-level amplification in CCNE2 was identified. (c) A patient was diagnosed with localized HR-/HER2+ breast cancer and treated with chemotherapy. Late metastatic relapse occurred with a new contralateral tumor, now HR+/HER2-. Following progression on tamoxifen, and prior to treatment with CDK4/6i and letrozole, an ERBB2 mutation was identified. Despite the presence of this alteration, the patient has had a durable ongoing response to CDK4/6i-based treatment.

**Supplemental Figure 4. Candidate alterations provoke CDK4/6i resistance *in vitro* (MCF7).**

(a) MCF7 cells were modified via CRISPR-mediated downregulation (RB1) or lentiviral overexpression (AKT1, KRAS G12D, AURKA, CCNE2) to interrogate potential resistance mediators identified in patient biopsy samples. Western blotting with the indicated antibodies is included. (b-f) Modified MCF7 cells were exposed to escalating doses of CDK4/6i (palbociclib – left, abemaciclib – right) and viability was estimated via cell-titer-glo (CTG) assay. Control (CRISPR non-targeting guide, GFP) cells are plotted along with the resistance driver of interest (RB1 – b, AKT1 – c, KRAS G12D – d, AURKA – e, CCNE2 – f). Parental and variant cell lines are normalized to vehicle control and viability is plotted as a function of increasing CDK4/6i (graphed as triplicate average +/-standard deviation). RB1, AKT1, and CCNE2 provoke CDK4/6i resistance (to both palbociclib and abemaciclib) *in vitro* in MCF7 cells. Corresponding IC50 values are included in Supplemental Table 7.

**Supplemental Figure 5. Candidate alterations provoke variable anti-estrogen resistance *in vitro*.**

Cell lines modified to reflect potential resistance drivers (per Figure 4 and Supplemental Figure 6; T47D – left, MCF7 - right) were exposed to escalating doses of fulvestrant (a – e). Drug response was assessed via cell-titer-glo (CTG) assay. Control (CRISPR non-targeting guide, GFP) cells are plotted along with the resistance driver of interest (RB1 – a, AKT1 – b, KRAS G12D – c, AURKA – d, CCNE2 – e). Parental and variant cell lines are normalized to vehicle control and viability is plotted as a function of increasing CDK4/6i (graphed as triplicate average +/- standard deviation). AKT1 and CCNE2 provoke fulvestrant resistance *in vitro* in both T47D and MCF7 cells. RB1 provokes minimal fulvestrant resistance in both T47D and MCF7. KRAS G12D and AURKA provoke significant fulvestrant resistance in T47D; KRAS G12D provokes minimal resistance in MCF7, while AURKA does not convey any resistance in MCF7. Corresponding IC50 values are included in Supplemental Table 7.

**Supplemental Figure 6. MDA-MB-361-AR-1 demonstrates upregulation of RAS-ERK pathway effectors via proteomic analysis.**

Digiwest proteomic analysis of MDA-MB-361-AR-1 cells versus parental MCF-7 cells demonstrates increased activation of multiple RAS-pathway effectors including KRAS, MEK, and ERK. These results suggest that the upregulation in pERK noted via western blot analysis correlates with pathway activation in the derivative cells.

## Supplementary Table Legends

**Supplemental Table 1. Clinical samples included in landscape analysis (excel file, 1 tab)**

Clinical information including treatment regimen, treatment duration (days), best radiographic response (BRR), and timing of the biopsy relative to treatment initiation/cessation (days). Biopsy sample information including receptor status, biopsy site, cancer-purity of sample and treatment-related information

**Supplemental Table 2. Clinical cohort characteristics (excel file, 1 tab)**

Clinical parameters of interest are included at the patient level (n = 58).

**Supplemental Table 3. Exome and mutational information (excel file, 3 tabs)**

Tab 1 – Exome-wide single nucleotide variants (SNVs) and Indels; Tab 2-Copy Number Variants (CNVs) at the segment level including Copy Number Above Ploidy (CNAP); Tab 3 - CNVs and Bi-Allelic inactivation at the single-gene level among oncogene and tumor suppressor gene candidates; Tab 4 – Genomic alterations among candidate mechanisms of resistance (MOR) among the resistance samples in our cohort. Candidate MOR genes include – RB1 with HOMDEL mutation type, AURKA - with Amplifications including GAIN) CCNE2 AKT1, RAS (KRAS, NRAS, and HRAS), ERBB2, and FGFR (FGFR1, FGFR2, and FGF3) – with activating evens – Amplifications and putative activating SNVs; Tab 5 – literature based list of known oncogenes (n=489) and tumor suppressor gene candidates (n=483).^45–48^

**Supplemental Table 4. Enrichment analysis of mutation in resistant vs. sensitive tumors (excel file, 1 tab)**

Fisher’s Exact test (single-side, for enrichment) comparing gene-specific the frequency of mutational events: HOMDEL==Bi-Allelic inactivation (among tumor suppressor candidates), IHC loss (for ER receptor), and gene activation by copy- number amplification – GAIN.up== CNAP>=1.5, AMP.up== CNAP>=3, or gene activation by either amplification or activating mutation – ACT==CNAP>=3 or Gain-of-function or recurring mutation, ACT.inc== same as ACT, but including non-recurring missense mutation (among oncogene candidates)

**Supplemental Table 5. Driver enrichment within patient populations (excel file, 1 tab)**

Sensitive, intrinsic resistant, and acquired resistant biopsies harboring any of the 8 potential driver alterations are quantified and graphed in figure 1D. Potential driver alterations include ER loss, amplification/mutation of ERBB2, FGFR2, CCNE2, AURKA, RAS, AKT1 and biallelic disruption of RB1.

**Supplemental Table 6. Evolutionary analysis and clonal fraction across 7 patients with multiple biopsies spanning pre- and post-treatment timepoints (excel file, 7 tabs)**

For each of the 7 patients with multiple biopsies, the clonal prevalence and evolutionary dynamic information is provided by depicting for each SNV (mutation_id) the cancer-cell fraction (cellular_prevalence) in each of the samples/time-point (sample_id), among other clone/cluster related information

**Supplemental Table 7. - IC50 Values for Drug Treatment Assays**

Corresponding IC50 estimates to the various drug response relationships provide in Figure 4 and Supplemental Figures 4 and 5 are provided here

**Supplemental Table 8. - IC50 Values for Culture to Resistance Experiments**

Corresponding IC50 estimates to the various drug response relationships provided in Figure 5 are provided here

