## Supplementary figures and images for "The genomic landscape of intrinsic and acquired resistance to cyclin-dependent kinase 4/6 inhibitors in patients with hormone receptor positive metastatic breast cancer"

### Supplemental Figure 1

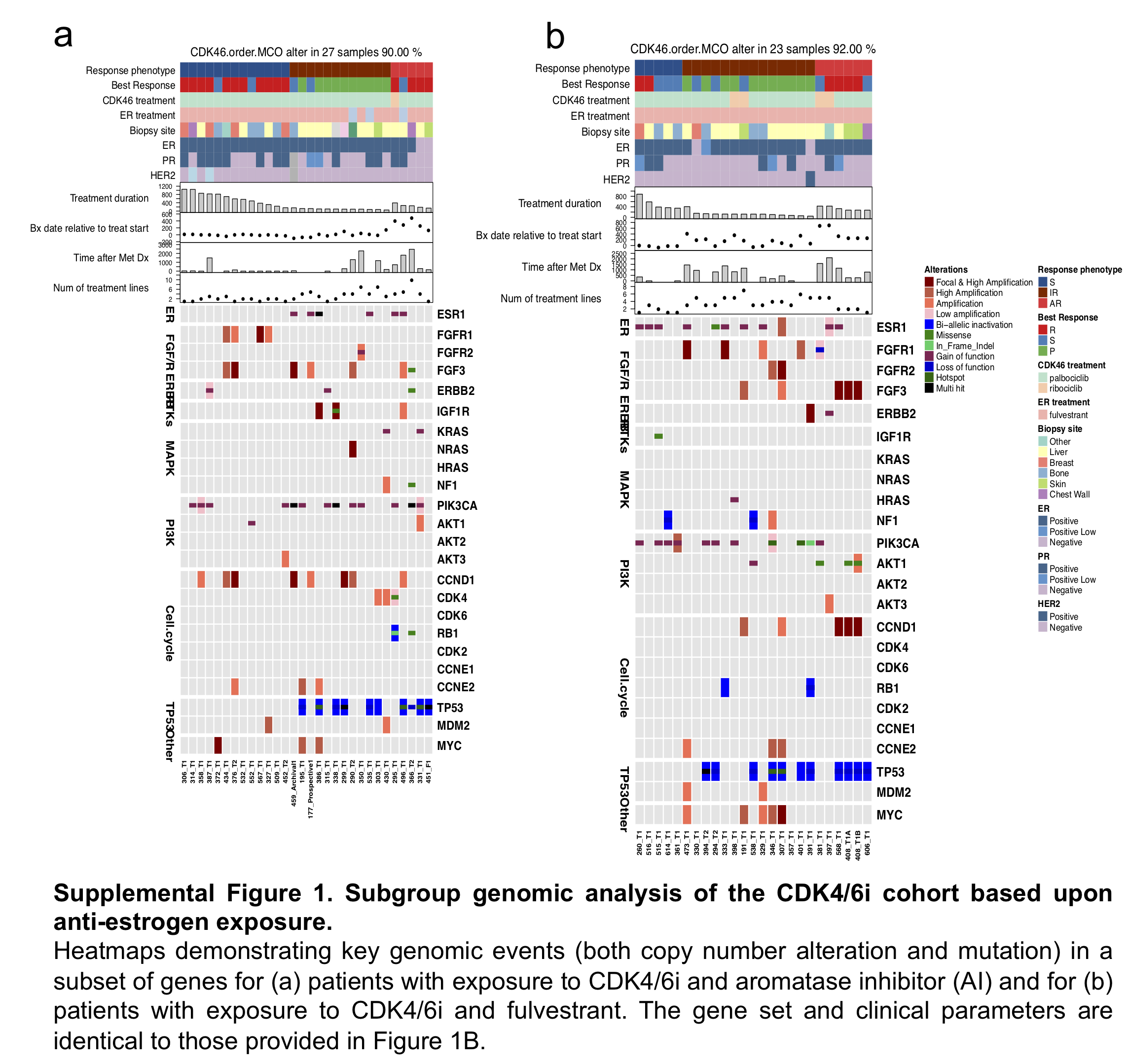

### Supplemental Figure 2

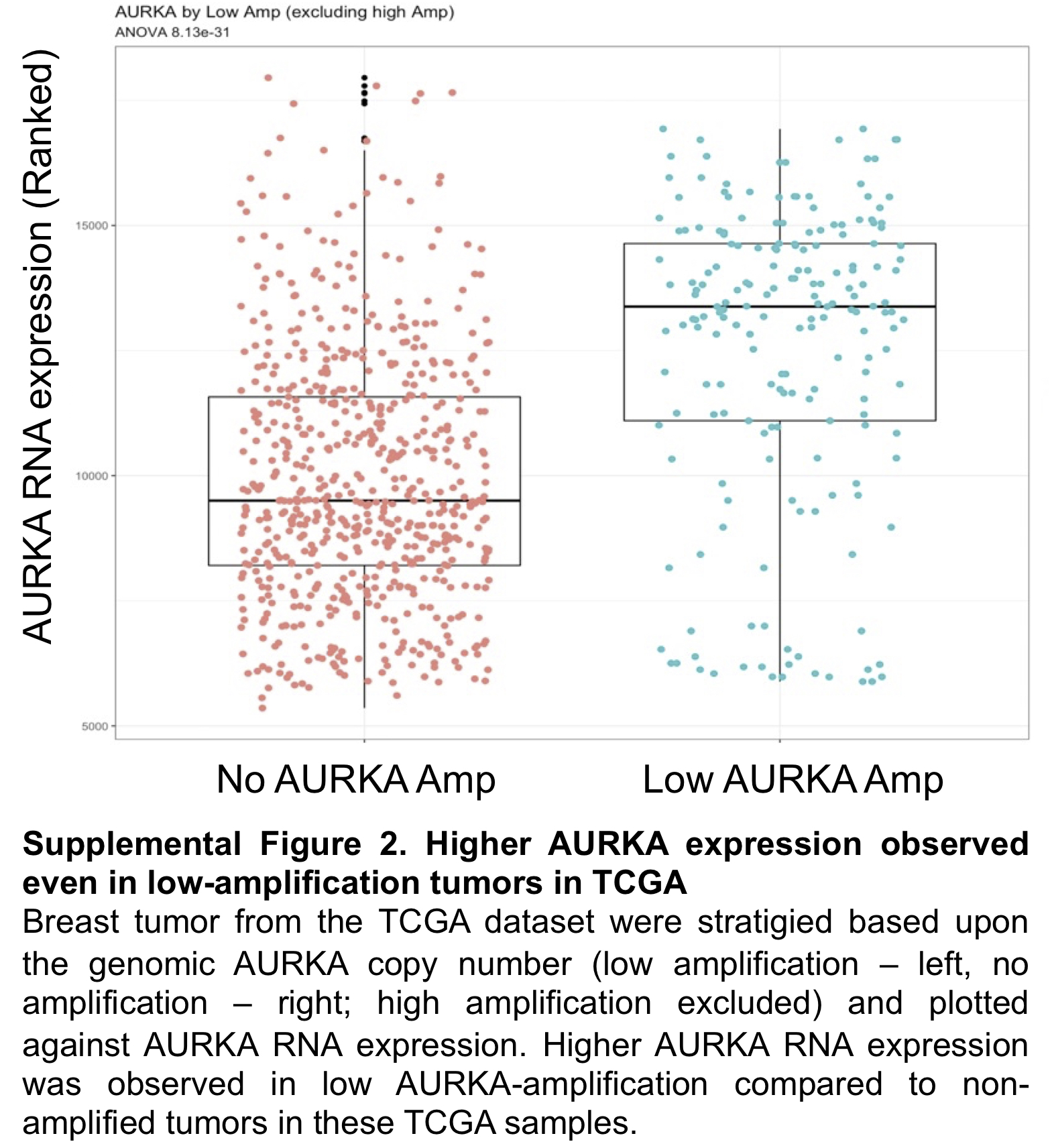

### Supplemental Figure 3

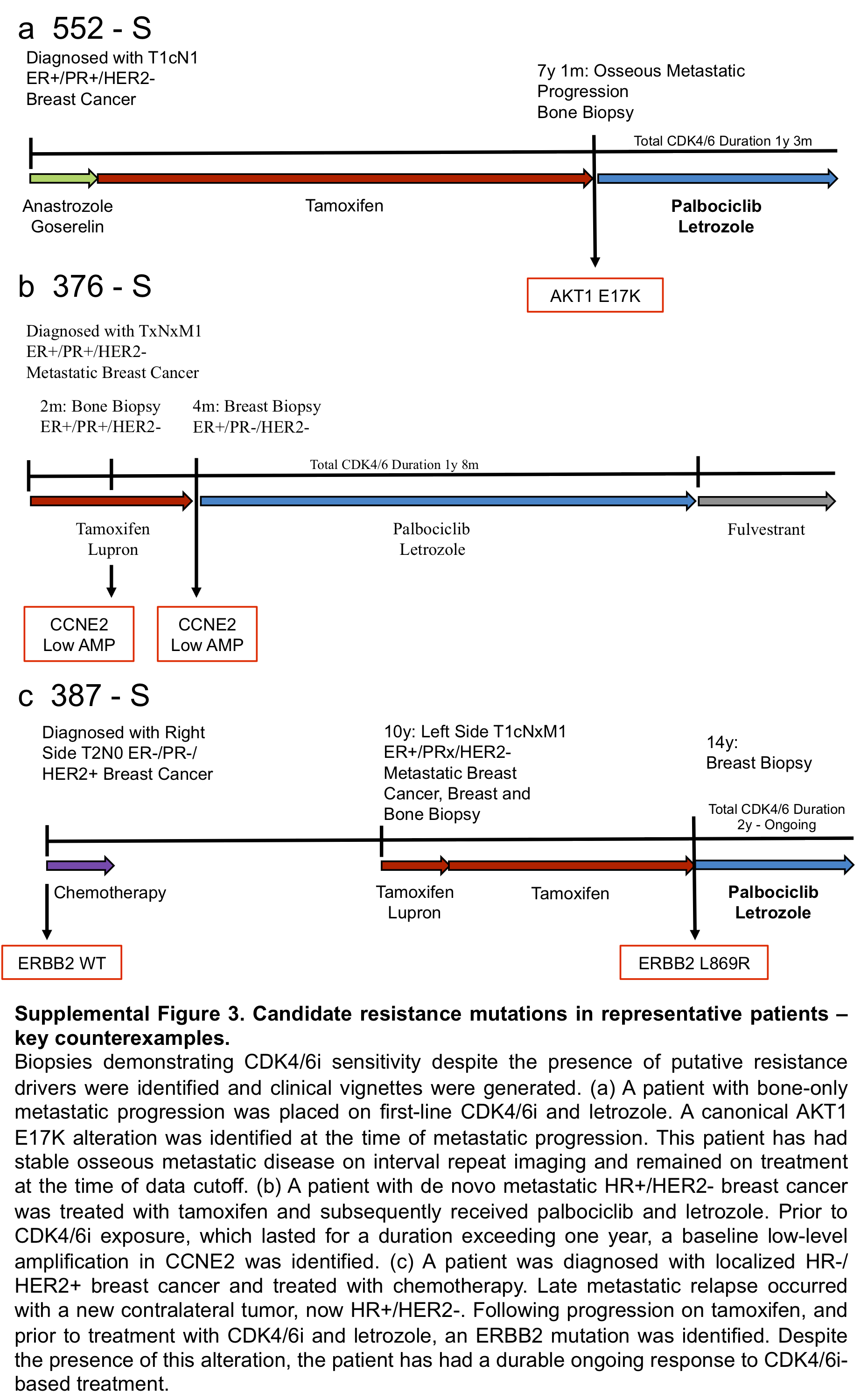

### Supplemental Figure 4

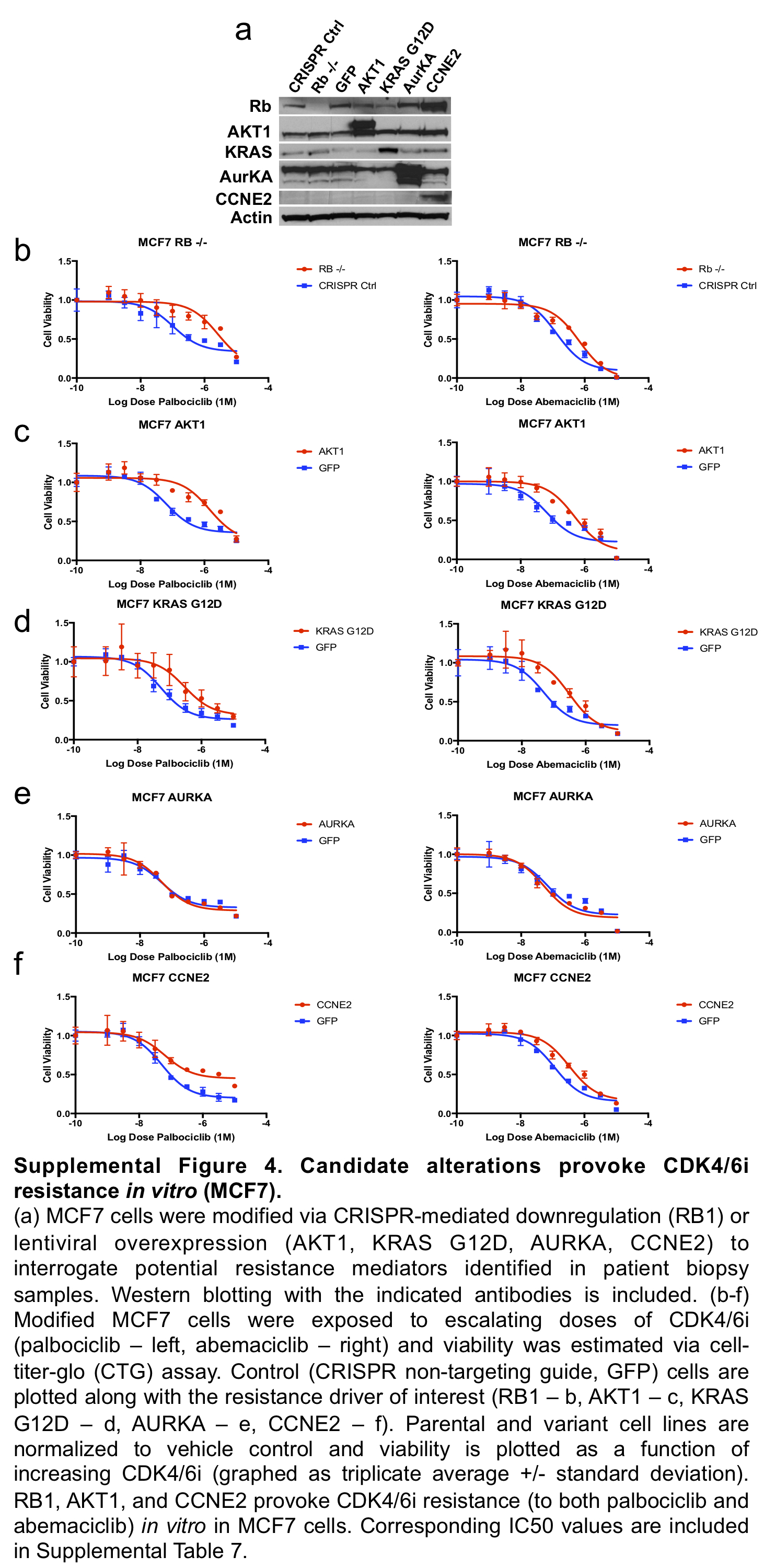

### Supplemental Figure 5

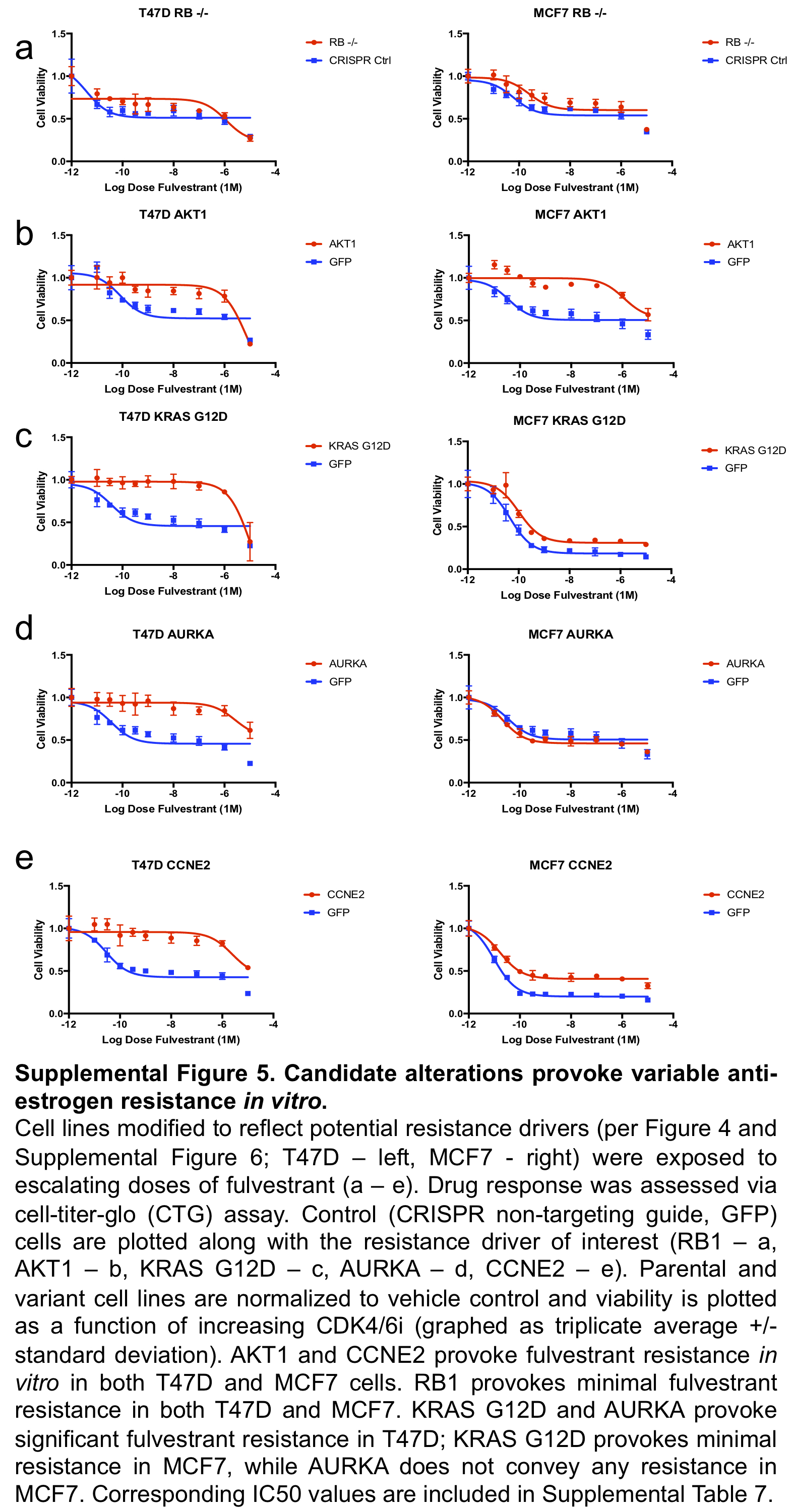

### Supplemental Figure 6

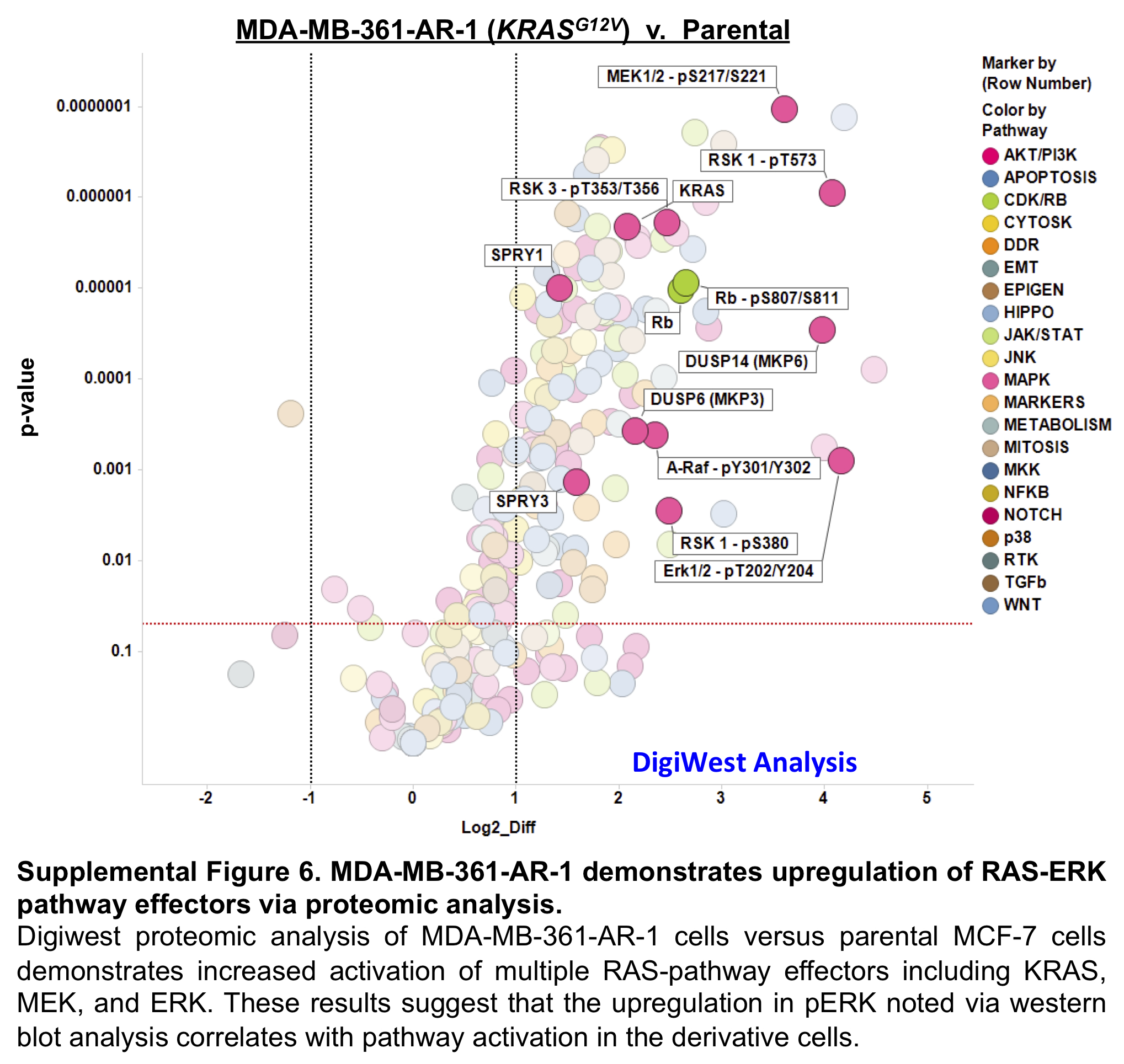
